# Design of antigens to present a tumor-specific cryptic epitope

**DOI:** 10.1101/2024.04.15.588917

**Authors:** Huafeng Xu, Timothy Palpant, Qi Wang, David E. Shaw

**Author notes:** To whom correspondence should be addressed. <u>Huafeng Xu</u>, David E. Shaw.

## Abstract

In many cancers, the epidermal growth factor receptor (EGFR) gene is amplified, mutated, or both. The monoclonal antibody mAb806 binds selectively to cancer cells that overexpress EGFR or express the truncated mutant EGFRvIII, but not to normal cells. This suggests that a promising avenue for developing cancer vaccines may be to design antigens that elicit mAb806-like antibodies. In this study, we designed antigens that present the mAb806-binding epitope in the same conformation as in overexpressed or truncated EGFR. We first used molecular dynamics simulations to identify conformations of EGFR in which the residues of the mAb806-binding epitope are accessible. We then designed antigens by substituting that epitope in place of a structurally similar loop in a different protein and generating mutants that could potentially stabilize the mAb806-binding conformation in this new context. Two mutants in which the epitope remained stable in subsequent simulations were chosen for evaluation in vitro. Binding kinetics experiments with these designed antigens provided strong evidence that the epitope was successfully stabilized in the mAb806-binding conformation, suggesting that they could potentially form the basis of vaccines that elicit cancer-selective antibodies.

## Introduction

Antigens—substances that induce antibody production—derive either from external sources, as occurs with infection or vaccination, or from endogenous sources, as in autoimmune disorders. Cancer cells are an endogenous source of antigens, and antibodies that target antigens on the surfaces of cancer cells have been used successfully as cancer therapies in a clinical setting.^1,2^ Current anti-cancer antibodies, however, are often cross-reactive against healthy tissues, which can lead to serious side effects.^3^ One avenue for improving specificity is to exploit a tendency for cancer cells to express surface proteins in abnormal conformations.^4,5^ These conformations may expose epitopes—the specific regions of antigens that are recognized by antibodies—that are sterically inaccessible in native protein conformations.^6^ (We will refer to such epitopes as cryptic epitopes.) Antibodies that can differentiate between native and abnormal conformations of the same protein have the potential to become a new class of therapeutics with a superior efficacy and safety profile.^7^ Antigens that can elicit such antibodies may both facilitate the development of therapeutic antibodies, and themselves be candidates for cancer vaccines that directly elicit immunological anti-cancer responses in patients.

Given a cryptic epitope and its three-dimensional structure in a non-native protein conformation, epitope-focused antigen design^8–14^ could potentially be used to develop antigens that present the epitope in the abnormal conformation and thereby elicit antibodies specifically recognizing that conformation. Non-native conformations, however, are often less stable—and thus rarer in the equilibrium population—than native conformations, and are thus often difficult to determine using experimental structural methods such as X-ray crystallography or nuclear magnetic resonance (NMR) spectroscopy. Molecular dynamics (MD) simulation^15^ could be a viable alternative for generating models of non-native conformations or other transient conformations that expose otherwise inaccessible epitopes.^16−18^

The mouse monoclonal antibody mAb806 binds selectively to cancer cells that either overexpress the epidermal growth factor receptor (EGFR) or express an exon 2–7 deletion mutant known as EGFRvIII.^19,20^ The 16-residue epitope recognized by mAb806 is accessible only in EGFR proteins that are misfolded as a consequence of overexpression, the exon 2–7 deletion,^21^ or some oncogenic mutations that alter the conformational dynamics of the EGFR ectodomain.^22^ A crystal structure of the complex between the mAb806 antigen-binding fragment (Fab) and its epitope peptide is available,^23^ offering one starting point for structure-based antigen design. The EGFR conformation to which mAb806 binds, however, remains unknown, and the epitope in mAb806-bound EGFR might have differences in conformation from that in the crystal structure of the peptide-mAb806 complex. Using MD simulations to model the conformation of the epitope in the context of full-length misfolded EGFR could thus increase the potential for designing a successful antigen.

In this study, we designed and tested antigens that present the EGFR cryptic epitope in its mAb806-binding conformation. We first used MD simulations to find conformations of EGFR in which the cryptic epitope was accessible for mAb806 binding. Starting from these conformations, we built structural models of the mAb806-EGFR complex. We then searched the Protein Data Bank for proteins with loop regions that are structurally similar to the epitope in the modeled mAb806-bound EGFR conformations, replaced the loop region of a candidate protein with the epitope, and designed a series of mutants with the aim of stabilizing the epitope in the mAb806-binding conformation. For many of the designed antigens, the epitope primarily occupied the antibody-binding conformation in subsequent MD simulations. From these antigens, we selected two for experimental characterization.

Binding kinetics experiments confirmed that these two designed antigens bound mAb806 with substantially higher association rate constants and binding affinities than the epitope peptide alone, providing strong evidence that the antigens successfully stabilized the epitope in the mAb806-binding conformation. In preliminary immunization experiments, our designed antigens were only weakly immunogenic, possibly due to their small size; nevertheless, their ability to present the epitope in the mAb806-binding conformation suggests they may be promising starting points for the development of a vaccine that elicits an immune response of mAb806-like, cancer-selective antibodies. More generally, our antigen design strategy, in which MD simulations generate models of target epitope conformations and are subsequently used to assess their stability in the context of designed antigens, may help facilitate the future development of highly selective, and thus safe, cancer vaccines.

## Results

To generate a structural model of mAb806 in complex with EGFR, we used long-timescale MD simulations, aided by an adaptive simulated tempering protocol (see Methods for details), to simulate the extracellular portion (ectodomain) of EGFR. The EGFR ectodomain consists of two ligand-binding domains (domains I and III) and two cysteine-rich domains (domains II and IV). In order to reduce the size of the molecular system and thus accelerate our simulations, we omitted domain IV, which is spatially remote from the epitope, and thus unlikely to affect its conformation or exposure. We performed two such simulations (referred to below as simulations 1 and 2), each starting from a different initial conformation drawn randomly from an equilibration simulation. We then attempted to dock the Fab of mAb806 onto the epitope in EGFR conformations taken from frames of our simulations by aligning the Cα atoms of the epitope in the crystal structure of the mAb806-epitope complex (PDB ID: 3G5V) with those of the epitope in the simulated EGFR conformations, and assessed for each conformation whether steric clashes occurred in the resulting mAb806-EGFR complex.

In its normal, regulated function, EGFR can transition from an auto-inhibited, tethered conformation to an active, untethered conformation in which domains I and II are rotated. Based on their X-ray crystal structures,^24,25^ steric hindrance would prevent mAb806 from binding to either of these conformations (Figure S1). It is possible that the epitope is exposed in an intermediate conformation during this transition,^6,23^ but this appears unlikely given that the specificity of the antibody indicates that it does not bind normally functioning wild-type (WT) EGFR. In simulation 1, EGFR transitioned between the tethered and untethered conformations while hiding the epitope from mAb806 (Figure 1, Movie S1), further supporting the hypothesis that mAb806 only binds to EGFR in aberrant conformations that are not necessary for its normal function.^21^

**Figure 1.**
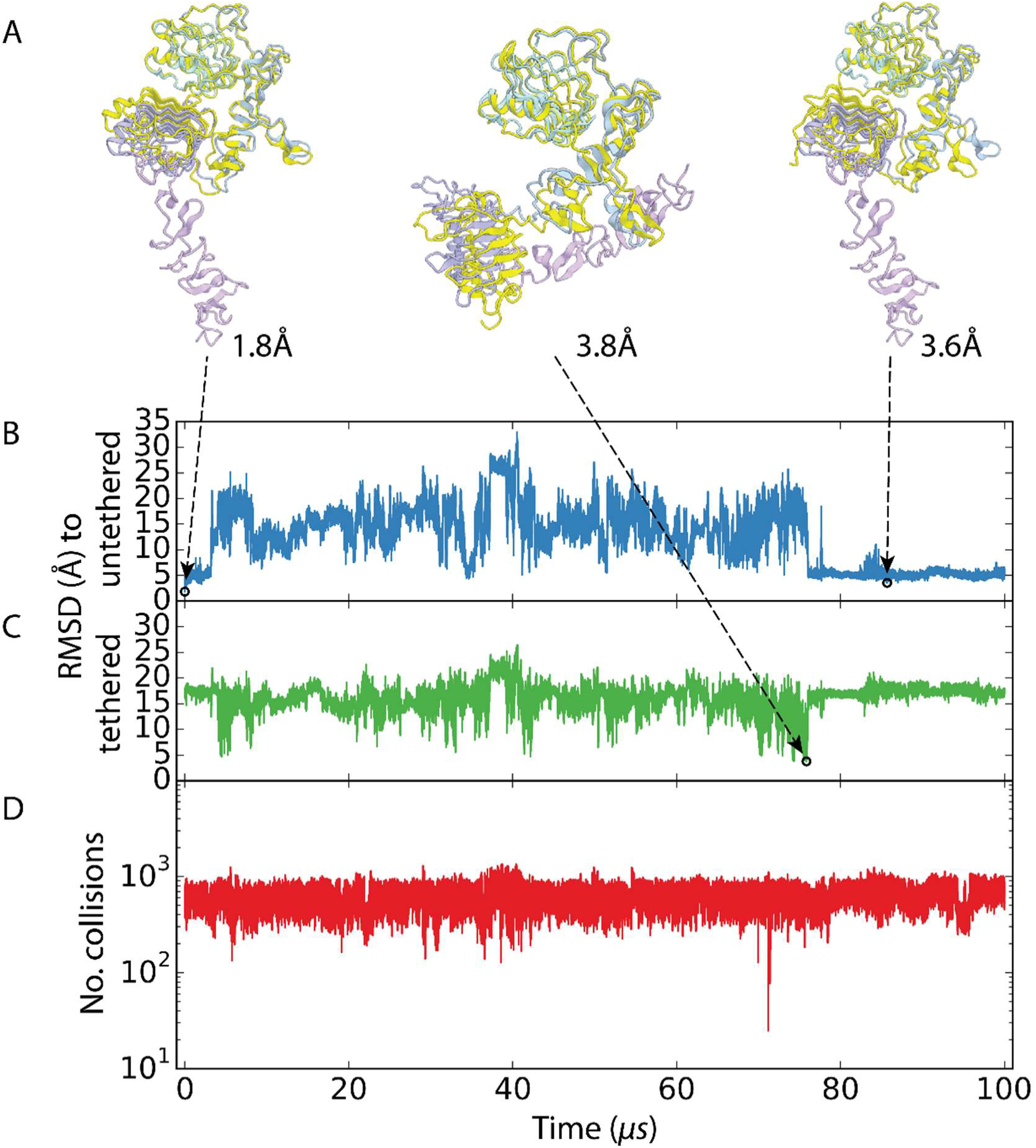
EGFR can transition between the untethered and tethered conformations without exposing the epitope to mAb806 binding. **A**) Snapshots of EGFR conformations from simulation 1 (using simulated tempering). Simulated conformations, shown in yellow, are superimposed onto crystal structures of the untethered conformation (left and right; PDB ID: 3NJP) or tethered conformation (center; PDB ID: 1NQL), in the crystal structures, domains I (also known as the L1 domain), II (the CR1 domain), III (the L2 domain), and IV (the CR2 domain) are shown in cyan, light blue, purple, and light purple, respectively. Domain IV was not included in our simulations. The simulation shown starts from an untethered conformation (left), visits a tethered conformation (center), and returns to an untethered conformation (right). The root-mean-square deviation (RMSD) of the Cα atoms of the conformations from the corresponding crystal structures are given beside the conformation. Also shown are time traces of the Cα RMSD from the crystal structures of **B**) the untethered conformation (blue) and **C**) the tethered conformation (green). **D**) The number of EGFR atoms that are in collision (interatomic distance ≤1.5 Å) with mAb806 docked onto its epitope remains greater than 20 in the simulation. mAb806 was docked onto each simulated conformation of EGFR by aligning the Cα atoms of the epitope in the crystal structure of the mAb806-epitope complex (PDB ID: 3G5V) onto the epitope in the simulated conformation.

In a small number of frames from simulation 2, the epitope, largely adopting the same conformation as in the crystal structure of the mAb806-epitope complex, became accessible to mAb806. When the mAb806 Fab was docked onto the epitope in these frames, there were no steric clashes between the Fab and EGFR (Figure 2, Movie S1). (A previous study simulating mAb806 binding to EGFR^26^ used steered MD simulation to promote exposure of the epitope to mAb806 binding.) This allowed us to build a structural model of mAb806 in complex with EGFR. We simulated the mAb806-EGFR complex starting from this structural model—once with simulated tempering (simulation 3) and once without (simulation 4)—and found that the complex remained stable in these simulations.

**Figure 2.**
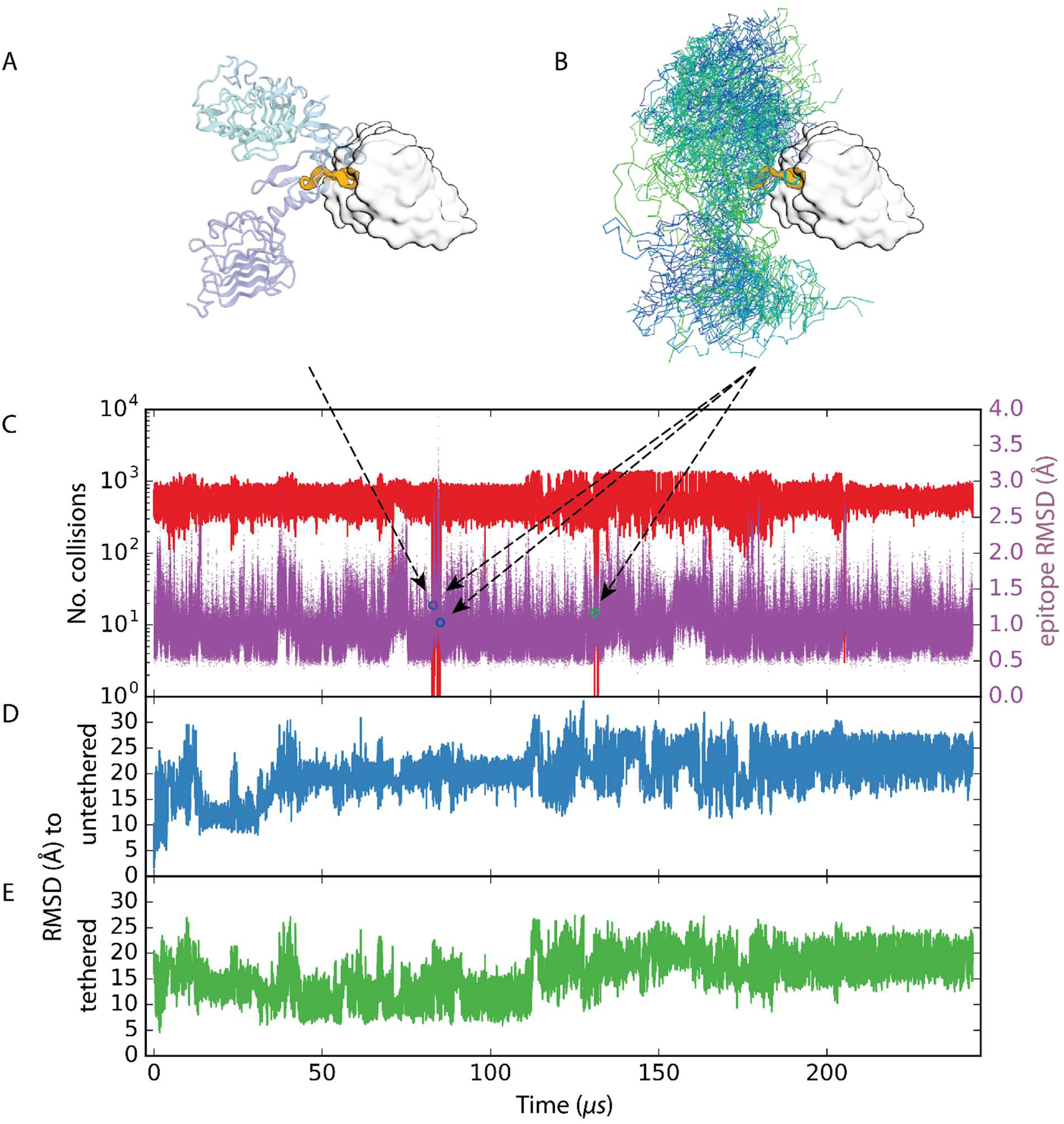
Local unfolding of EGFR permits mAb806 binding. We considered an EGFR conformation to be binding-competent if it exhibited no steric clashes with a docked mAb806, and the Cα RMSD of the epitope from its conformation in the crystal structures of the mAb806-epitope complex was no greater than 1.5 Å. **A**) Example of a binding-competent conformation sampled from simulation 2 (using simulated tempering). EGFR is shown as a ribbon diagram, with its I, II, III domains colored as in Figure 1A. The epitope is highlighted as orange. The docked mAb806 Fab is shown by its molecular surface, in white with black contours. **B**) Multiple locally unfolded, binding-competent EGFR conformations (shown in backbone trace, colored serially in gradient from blue to green) are sampled in the same MD simulation. **C**) The number of EGFR atoms in collision with a docked mAb806 (red) and the Cα RMSD of the epitope from its binding conformation (purple) in the simulation. The binding-competent conformations are highlighted by circles colored serially as in panel (B). Also shown are the Cα RMSD from the crystal structures of **D**) the untethered conformation and **E**) the tethered conformation in the simulation. The binding-competent conformations are different from both the tethered and untethered conformations.

The mAb806-binding conformation from our MD simulations differs from native EGFR conformations by a rotation in the Cα–C bond of the Arg285 residue; that is, a change in the ψ dihedral angle (Figure 3). This flips the epitope such that it no longer contacts the adjacent β-sheet (residues Try275 to Arg285) and exposes it for mAb806 binding. Mutationally disrupting this β-sheet destabilizes its contacts with the epitope, making the epitope more accessible to mAb806.^23^ In agreement with this finding, we found that the mutant EGFR^C271A/C283A^, which eliminates a disulfide bond that restrains the β-sheet’s conformation, was more prone to exposing the epitope to mAb806 binding in our simulations (simulations 5 and 6; Figure S2). These results offer a plausible explanation for mAb806’s selectivity for cells overexpressing EGFR or expressing EGFRvIII: In cells overexpressing EGFR, inadequate chaperone assistance might cause incorrect disulfide bonding patterns in domain II (also known as the CR1 domain), and in cells expressing EGFRvIII, the deletion—of residues 6 to 273—leads to the loss of the disulfide bond between Cys271 and Cys283, potentially disrupting the Tyr275–Arg285 β-sheet and thus exposing the epitope to mAb806 binding.

**Figure 3.**
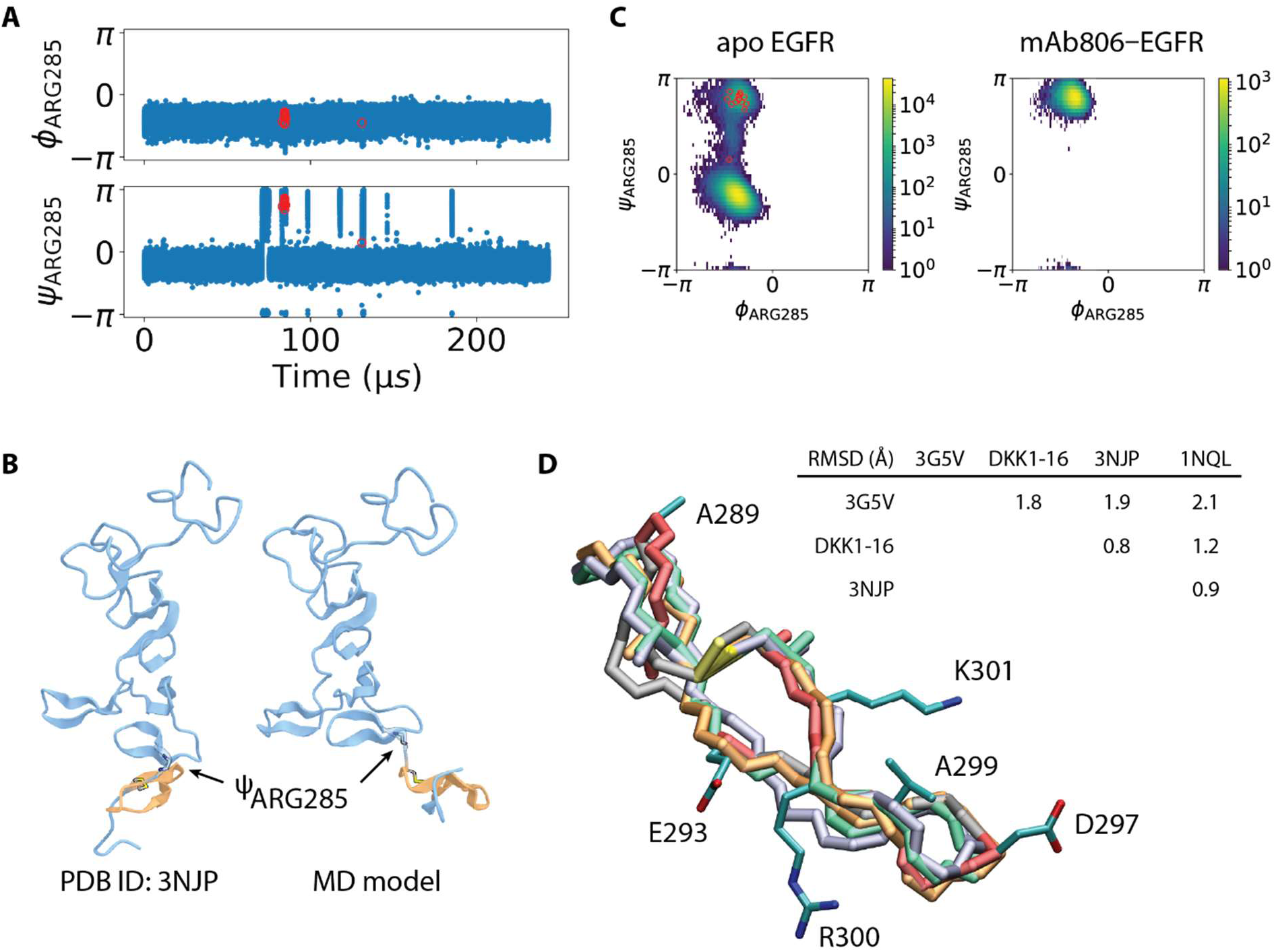
Computational model of the mAb806-binding conformation of EGFR. **A**) Time trace of the backbone ϕ and ψ dihedral angles of residue Arg285 in simulation 2. The snapshots in which the mAb806 Fab could be docked onto the epitope without steric clash are indicated by red circles. The epitope was only accessible when the ψ dihedral departed from its value in the native conformations. **B**) Domain II in the native, untethered conformation, and in the mAb806-accessible conformation generated in simulation 2. In the mAb806-binding conformation, the epitope (orange) flips around Arg285 and breaks its contact with the preceding β-sheet (Tyr275 to Arg285), exposing itself. **C**) The Ramachandran plots of the backbone ϕ,ψ dihedral angles of Arg285 in the simulation of the apo EGFR ectodomain (simulation 1) and in the simulation of the mAb806 Fab-EGFR complex (simulation 5). The bound Fab prevents Arg285 from flipping back to the native conformation. **D**) The mAb806-epitope in crystallographic and modeled conformations. The backbone traces of different conformations are colored as follows: the untethered conformation (PDB ID: 3NJP) in light green, the tethered conformation (PDB ID: 1NQL) in ice blue, the epitope peptide in complex with mAb806 (PDB ID: 3G5V) in gray, with the residues in contact with mAb806 highlighted in pink, and the epitope in the designed DKK1-16 in orange. The epitope structure in DKK1-16 is taken from a snapshot in the simulation of the mAb806 Fab–EGFR complex (simulation 4), except for the side chains of the redesigned residues. Sidechains in contact with mAb806 are shown as sticks. The RMSD between Cα atoms in the different conformations are shown in the table. The structure of the 16-mer epitope-peptide bound to mAb806 has notable differences from the structures of the epitope in EGFR.

The free 16-mer epitope peptide (sequence CGADSYEMEEDGVRKC, with a disulfide bond between the cysteines at positions 1 and 16, making it a cyclic peptide) is unstructured, and did not retain its mAb806-binding conformation in our simulations (Figures S3A and 4E), in agreement with a previous NMR experiment.^23^ This may limit the ability of the epitope peptide to function as an antigen to elicit mAb806-like antibodies. To design antigens that would present the epitope in the mAb806-binding conformation, we used as a starting point the ensemble of conformations generated from our MD simulation of the mAb806-EGFR complex. Using MotifGraft in Rosetta,^12,27^ we searched a subset of PDB structures for scaffolds with an appropriate segment; that is, a segment that could be replaced with the epitope in a conformation taken from a snapshot of our mAb806-EGFR simulation (see SI for details).

This search identified the cysteine-rich, 93-residue C-terminal domain of the human DKK1 protein (PDB ID: 3S2K, chain C) as a potential scaffold. In four simulated conformations, the 16-residue epitope could be superimposed onto a 16-residue hairpin of this scaffold protein without substantial steric clashes occurring between the Fab and the scaffold protein (Figure 4). In contrast, using the crystal structure of mAb806-epitope complex would not have identified this scaffold (Figure 4D). This suggests that, for antigen design in general, including a conformational ensemble of an epitope from MD simulations can increase the chance of successfully identifying a suitable grafting scaffold.

**Figure 4.**
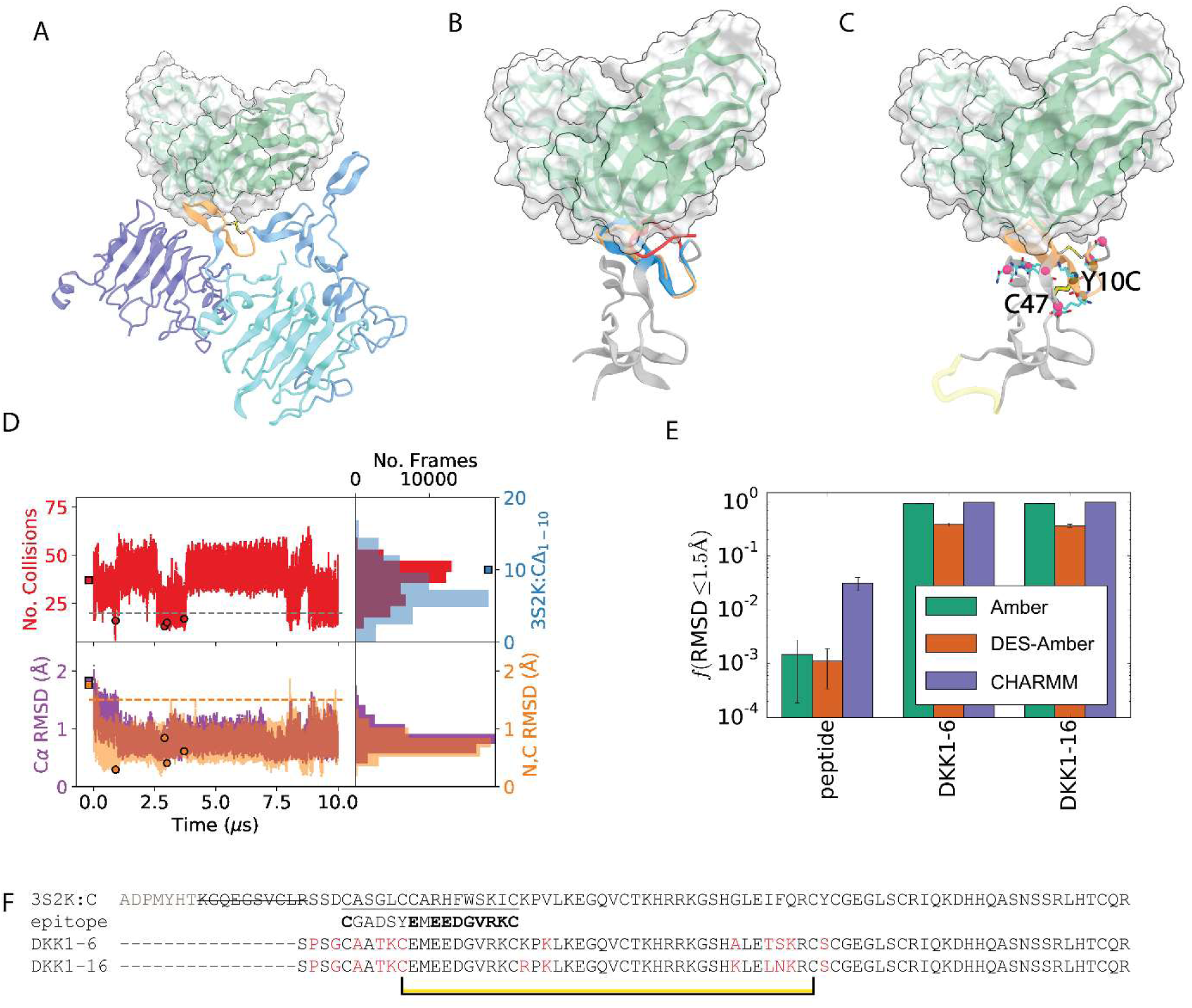
The design of antigens. **A**) Model of the mAb806-EGFR ectodomain complex, generated from an MD simulation, used as the starting point for antigen design. The 16-residue epitope is highlighted in orange, while domains I, II, and III of EGFR are shown as a ribbon diagram and colored as in Figure 1A. The Fab of mAb806 is shown as a ribbon diagram overlaid with its molecular surface. **B**) The C-terminal domain of human DKK1 (PDB ID: 3S2K, chain C), shown as a ribbon diagram and colored gray, was identified as a potential scaffold, allowing four epitope conformations—taken from an MD simulation of the mAb806-EGFR complex, such as that in (A)—to be superimposed onto the hairpin structure of residues C14–C29 (blue). Visual inspection suggests that the first ten N-terminal residues in the scaffold, shown in red, sterically clash with the antibody and should be deleted from the graft. **C**) Visual inspection suggests that the mutation Y10C may introduce a disulfide bond, shown as yellow sticks, with Cys47. RosettaDesign was used to identify additional mutations that might stabilize the conformation of the designed antigen. The residues mutated in the final design of DKK1-16 are highlighted by purple spheres. The loop that was missing in the crystal structure but modeled by Rosetta is shown in yellow. **D**) The design was enabled by the MD simulation, which yielded EGFR-mAb806 Fab conformations in which the epitope could be grafted into the scaffold protein DKK1. The top panel shows the steric clashes between the scaffold protein DKK1 with the epitope-mAb806 complex, when the latter is superimposed onto the former by aligning the Cα atoms of the epitope onto the Cα atoms of the replacement segment C14–C29. The time trace is shown on the left, and corresponding histogram on the right. The light blue bars in the histogram on the right show the number of atoms that would clash if the first ten N-terminal residues in DKK1 were removed (right y-axis). The bottom panel shows the Cα RMSD (purple, left y-axis) and the N, C, and Cα RMSD values (orange, right y-axis) of the N- and C-terminal residues between the epitope and the replacement segment. The conformational snapshots that were successfully grafted onto the DKK1 scaffold protein are highlighted in circles. The values corresponding to the crystal structure of the epitope-mAb806 complex (PDB ID: 3G5V) are shown as squares; a search using the epitope-mAb806 conformation from the crystal structure did not identify DKK1 as a potential candidate. **E**) In our MD simulations of the designed antigens DKK1-6 and DKK1-16 using three different force fields (Amber: Amber99SB*-ILDN + TIP3P water model; DES-Amber: DES-Amber SF1.0 + TIP4P-D water model; CHARMM: CHARM22* + TIP3P-CHARMM water model), the epitope mostly occupies the mAb806-binding conformation (Cα RMSD from the crystal structure ≤1.5 Å), whereas the free epitope peptide (cyclized due to the C1–C16 disulfide bond) rarely visits the binding conformation. The error bars were generated by dividing each simulation into five non-overlapping time intervals and computing the standard deviations between those intervals. **F**) Amino acid sequences of the two designed antigens DKK1-6 and DKK1-16 aligned to that of the scaffold DKK1 protein and the epitope C287–C302 in EGFR. The gray letters in the N-terminus of the scaffold protein indicate the unresolved residues in the crystal structure (PDB ID: 3S2K), and the struck-through letters signify the ten residues in the scaffold protein that were deleted in the designed antigens to avoid steric clashes with mAb806. The bold-faced residues in the epitope form specific interactions with mAb806 (except for the two terminal cysteines, which form the critical disulfide bond), and were not considered for mutation in the remodeling step. The residues mutated in the remodeling step are highlighted in red. The connection between Cys10 and Cys47 signifies the manually designed disulfide bond.

Visual inspection of the epitope-grafted scaffold led us to introduce an additional disulfide bond by changing the Tyr at residue 10 of the antigen to Cys (Figure 4C). Subsequently, 17 mutants were computationally generated using RosettaRemodel^28^ in order to resolve steric clashes and form new specific interactions that would stabilize the designed protein conformation (Figure 4C). For each designed mutant, MD simulations totaling 10 to 100 μs of simulation time were carried out using three different force fields (Methods). For the majority of the mutants, the epitope retained or predominantly adopted the mAb806-binding conformation in our simulations (Figure S3B). After visual inspection of the structures of the designs, we chose to pursue two, DKK1-6 and DKK1-16 (Figure 4F), for additional experimentation.

Both designed proteins expressed well in Sf21 cells. The purified proteins were characterized by 1D proton NMR and showed signatures indicating folded proteins (Figure S4A). The similarity between the spectra of DKK1-6 and DKK1-16 suggests that the two proteins adopt the same fold (Figure S4A). The proteins were induced to unfold by the reduction of their disulfide bonds, suggesting that those disulfide bonds are important in maintaining the proteins’ correct fold (Figure S4B).

We characterized the binding of the designed antigens against ch806, a chimeric version of mAb806 with a human fragment crystallizable (Fc) region. Both DKK1-6 and DKK1-16 had substantially higher association rate constants for binding to ch806 than the free epitope peptide [Cyclo(1,16)] H_2_N-CGADSYEMEEDGVRKC-OH (Figures 5A and S5). Their association rate constants (>10^6^ M^−1^ s^−1^) are in the range for diffusion-limited antibody-antigen binding,^29^ suggesting that the epitope in the designed antigens is stabilized in the ch806-binding conformation,^30^ consistent with our MD simulation results (Figure 4E, Figure 5A inset). The increased binding affinity of the designed antigens over the epitope peptide is demonstrated by both surface and solution measurements (Figure 5B). The antigen DKK1-16 is also shown to form a stable complex with the mAb806 Fab (Figures 5C and 5D). Taken together, these results suggest that the designed antigens likely stabilize the epitope in the mAb806-binding conformation.

**Figure 5.**
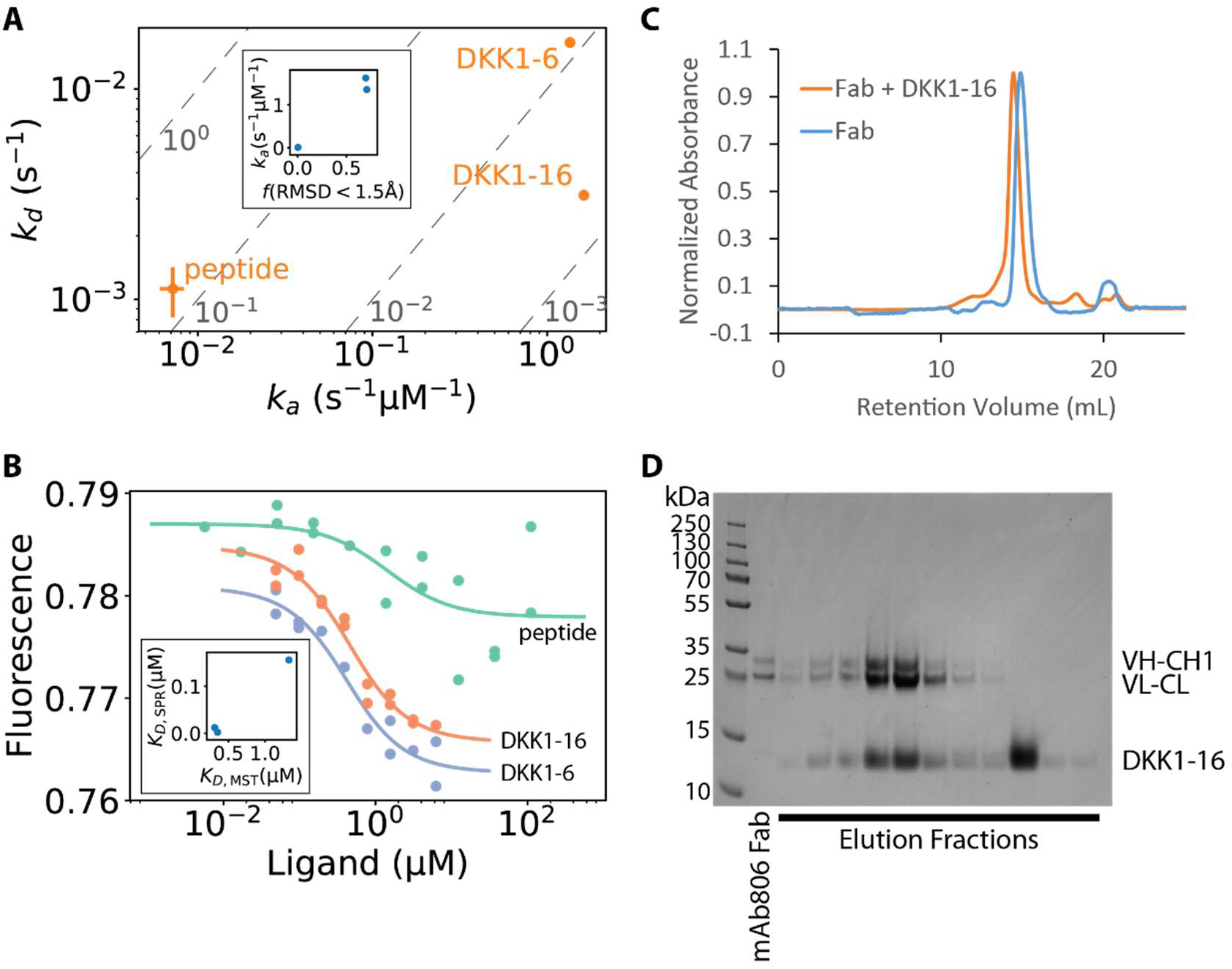
Experimental characterization of the binding of designed antigens to the ch806 antibody. **A**) Surface plasmon resonance measurements show that the designed antigens DKK1-6 and DKK1-16 bind to ch806 orders-of-magnitude faster than the free epitope peptide, suggesting that the epitopes in the designed antigens have been stabilized in the binding-competent conformation. The binding affinities are constant along the dashed diagonal lines, with the corresponding equilibrium dissociation constants, *K_D_*, indicated in gray. The high conformational stability of the epitope in the designed antigens, as estimated from MD simulations (Fig 4E), is consistent with the fast association rate constants of mAb806 binding determined by SPR (inset). **B**) Microscale thermophoresis (MST) measurements of the ch806-binding affinities for the epitope peptide (green, *K_D_* = 1.33 μM), DKK1-6 (blue, *K_D_* = 0.31 μM), and DKK1-16 (orange, *K_D_* = 0.35 μM). The inset compares the *K_D_* values determined by SPR and MST. MST confirmed that the designed antigens have higher affinity than the free epitope peptide for ch806 in bulk solution, suggesting that the improved binding measured by SPR is not an artifact of binding to the surface. **C**) Co-elution of mAb806 Fab and DKK1-16 in analytical size-exclusion chromatography. The association between the Fab and the designed antigen led to a shift in the retention volume that corresponds to a difference in molecular weight of ∼10 kDa, in agreement with the molecular weight of DKK1-16. **D**) The Fab and designed antigen in the co-elution peak were confirmed by SDS-PAGE. VH-CH1 and VL-CL are the heavy and the light chains, respectively, of the Fab. These results demonstrate that the designed antigen indeed forms a stable complex with mAb806 in solution.

We next tested the immunogenicity of the designed antigens. Rabbits were immunized with DKK1-6, DKK1-16, and the epitope peptide ([Cyclo(1,16)] H_2_N-CGADSYEMEEDGVRKCGGGS(KAoa)-amide) conjugated to keyhole limpet hemocynanin (KLH), a carrier protein (Figure 6A). We assessed both the anti-sera and the affinity-purified polyclonal antibodies (pAbs). As expected, based on their small molecular sizes, the designed antigens were on average much less immunogenic than the peptide-KLH conjugate in our enzyme-linked immunosorbent assay (ELISA) (Figures 6B and 6C). The pAbs from rabbit 2842, immunized with peptide-KLH conjugate and affinity purified using the epitope peptide, showed high affinity for the epitope peptide. In contrast, the pAbs elicited by, and affinity-purified using, the designed antigens showed much weaker affinity for the epitope peptide (Figure 6D). This might be partly because only a small fraction of the pAbs were targeting the epitope, while the majority of the pAbs were targeting other parts of the antigen.

**Figure 6.**
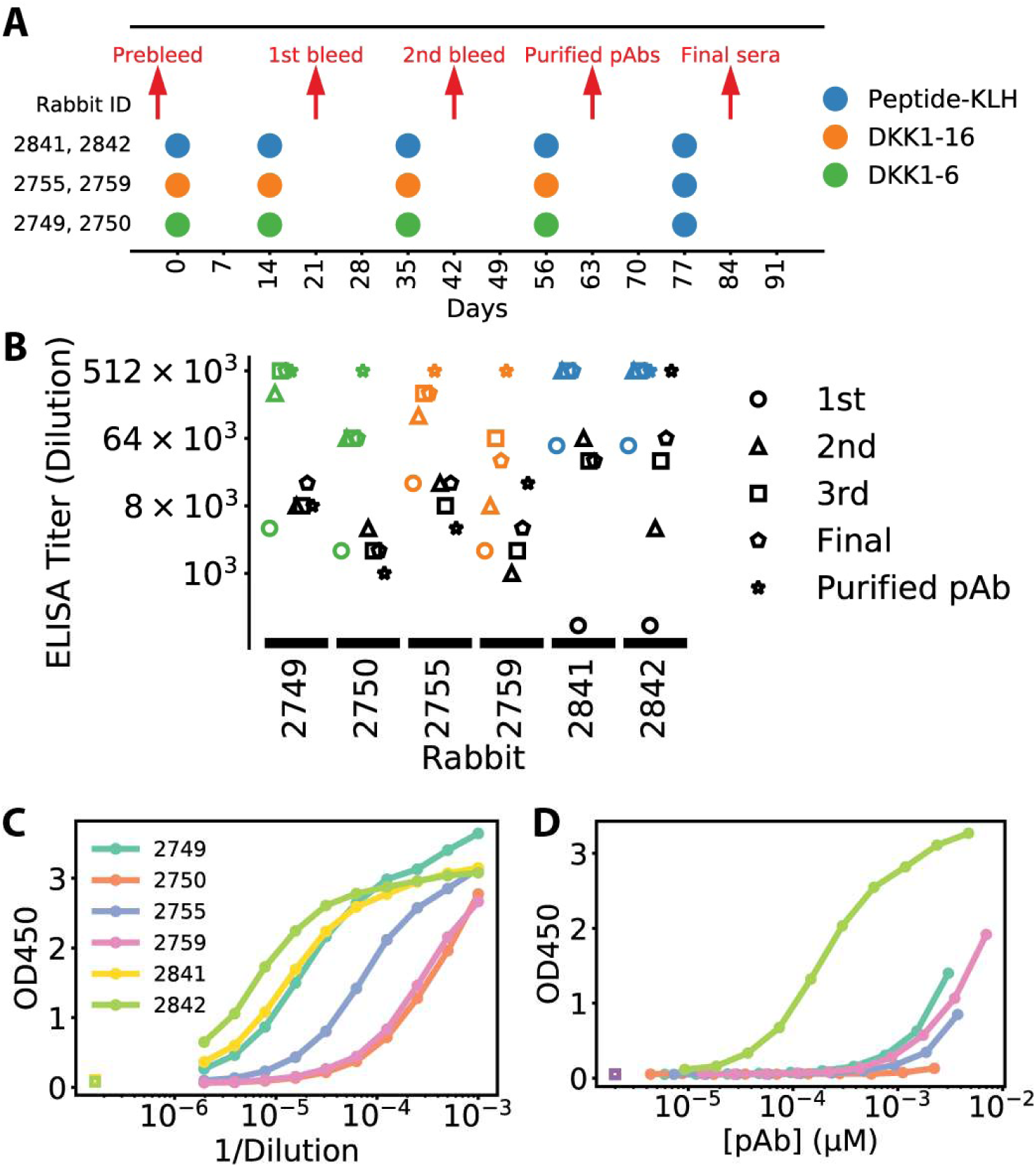
The designed antigens are capable of eliciting antibodies that bind to the epitope peptide. **A**) Rabbit immunization protocol. Two rabbits were immunized with each antigen on the indicated days (rabbits 2749 and 2750 received DKK1-6, rabbits 2755 and 2759 received DKK1-16, and rabbits 2841 and 2842 received peptide-KLH). Pre-immunization sera (prebleed) were obtained three days prior to the first immunization, and anti-sera were obtained on the indicated days. Polyclonal antibodies were affinity-purified from the third bleed. All rabbits received peptide-KLH as the final boost to stimulate antibody response against the mAb806 epitope. **B**) ELISA analysis of antigen and epitope binding of the antisera. The colored markers indicate the titers of binding to the eliciting antigen, whereas the black markers indicate the titers of binding to the epitope peptide [Cyclo(5,20)] H_2_N-SGGGCGADSYEMEEDGVRKC-OH. The final antisera and the purified pAbs from the rabbits immunized with the designed antigens all showed clear but weak binding to the epitope peptide, compared to much stronger binding to the eliciting antigen. One possible explanation is that only a small fraction of the antibodies from the humoral response were targeting the mAb806 epitope. **C**) ELISA readout of binding to the eliciting antigens at different dilutions of the third bleed sera, from which the pAbs were purified. The humoral immune response to the peptide-KLH immunogen was robust in both rabbits, whereas the response to the designed antigens tended to be weak (with the exception of rabbit 2749), likely a reflection of the poor immunogenicity of the designed antigens due to their small molecular sizes. **D**) ELISA readout of binding to the epitope peptide at different concentrations of the purified polyclonal antibodies. The pAbs elicited by the epitope-peptide KLH-conjugate and affinity purified by the epitope peptide bound to the epitope peptide with high affinity, whereas those elicited and affinity purified by the designed antigens had much lower affinity for the epitope peptide, suggesting that only a small fraction of the latter pAbs, which could bind to any part of the designed antigen, were binding to the epitope peptide.

We next tested how strongly the anti-sera and pAbs could bind to a set of five different tumor cell lines, including two that express normal levels of EGFR (U87MG and A549), two that overexpress EGFR (A431 and U87MG^wt^ ^EGFR^; U87MG^wt^ ^EGFR^ is a stable engineered U87MG cell line overexpressing EGFR, Figure S6), and a stable engineered U87MG cell line expressing EGFRvIII (U87MG^EGFRvIII^, Figure S6). We validated the EGFR profile of these cells by measuring ch806 binding to them: Both fluorescence activated cell sorting (FACS) and On-Cell Western assays showed that ch806 selectively binds to the latter three cell lines, with undetectable binding to the first two (Figures 7A and S7A). Quantitative analysis of the On-Cell Western titration curves (Figure S7B) for ch806 and cetuximab,^31^ a monoclonal antibody that binds to normal EGFR, suggests that ch806 binds to a subpopulation of EGFR on the cell surface (Figure S7C). These results are in agreement with previous experimental findings.^19,21^

**Figure 7.**
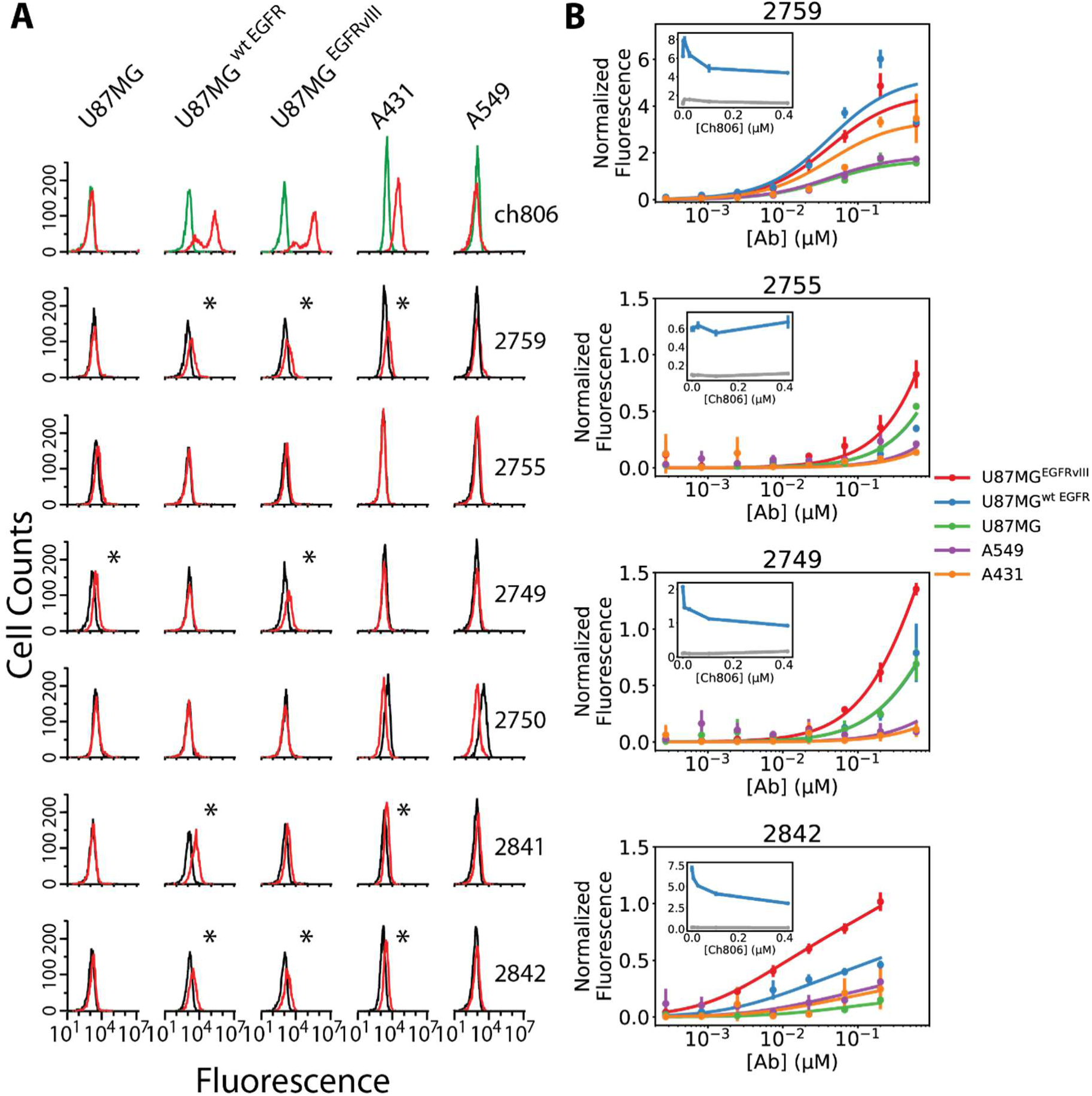
Rabbit polyclonal antibodies elicited by the epitope-peptide keyhole limpet hemocyanin (KLH) conjugate and the designed antigens bound weakly, and in some cases not at all, to tumor cells. **A**) FACS analysis of the final antisera binding to tumor cell lines. None of the antisera exhibited as high a binding affinity to any of the cell lines as the monoclonal ch806 binding to the three cell lines U87MG^wt^ ^EGFR^, U87MG^EGFRvIII^, and A431. Nevertheless, small shifts in the fluorescence distributions—indicative of some binding—were observed for some pairs of cell lines and antisera, as indicated by the asterisks. In particular, the antisera from rabbit 2759 and 2842 exhibited binding to the same three cell lines as ch806. The red lines are the binding data for the monoclonal antibody ch806 or the antisera, the green lines are those for human IgG1 control (Biolegend, Catalog No. 403102), and the black lines are those for the pre-immunization bleed of the corresponding animals. **B**) Characterization of the binding of the purified polyclonal antibodies to the tumor cell lines by an On-Cell Western assay. All pAbs demonstrated some binding to the cell line expressing EGFRvIII mutant, suggesting that they targeted an epitope that was more exposed in this mutant. All pAbs elicited by designed antigens, however, showed binding only at high concentrations, consistent with their low ELISA titers against the epitope peptide. Insets (in which the y-axis is also normalized fluorescence): Ch806 was able to compete against the binding of the polyclonal antibodies from rabbits 2759, 2749, and 2842 to U87MG^wt^ ^EGFR^ cells, suggesting that these polyclonal antibodies contain antibody molecules that bind to the same EGFR epitope on the tumor cells as the ch806 antibody. The rabbit pAb concentrations in these competition experiments were 67 nM (blue) and 0 nM (gray). The inhibition of binding by the ch806 antibody was only partial even at high ch806 concentrations, suggesting that the pAbs may have recognized a partial segment of the ch806-epitope that is inaccessible in normal EGFR conformations.

All the anti-sera and the pAbs from our immunizations bound to the A431, U87MG^wt^ ^EGFR^, and U87MG^EGFRvIII^ cells with much weaker affinity than did ch806 (Figure 7). The weak binding of the anti-sera and pAbs from rabbit 2842 to these cells contrasts with their high affinities for the epitope peptide, suggesting that the pAbs elicited by the unstructured epitope peptide do not recognize the mAb806-binding conformation of the epitope in EGFR. Notably, the weak binding of all the pAbs to U87MG^wt^ ^EGFR^ was only partially inhibited by ch806 (Figure 7B insets), which suggests that a subpopulation of the antibodies might have bound to a partial epitope that is exposed in normal EGFR, and that their binding did not compete with that of ch806, which primarily binds to misfolded EGFR. (It is unlikely that pAbs elicited by the designed antigens bound to U87MG cells because of antibodies reactive against DKK1, as U87MG has been shown to have undetectable expression of the *dkk1* gene^32^). Nevertheless, our finding that ch806 partially inhibits the binding of the pAbs indicates that at least a subpopulation of the antibodies is competing with ch806 for the same epitope.

## Discussion

Our MD simulations of EGFR in complex with the antibody mAb806 have suggested a plausible, atomistically detailed model of how mAb806 selectively binds to misfolded EGFR, and enabled us to design antigens that present the mAb806-recognizing epitope in its antibody-binding conformation. Structural stabilization of such antigens might be necessary in order to elicit mAb806-like antibodies, as the epitope peptide on its own is unstructured. This is supported by the results of our preliminary immunization experiment, in which antibodies elicited by the epitope peptide had high affinity for the peptide, but only weakly bound to overexpressed EGFR or EGFRvIII on tumor cell lines. Despite being stabilized, our designed antigens, DKK1-6 and DKK1-16, appeared to suffer from weak immunogenicity and the inability to focus the antibody response against the target epitope. Nonetheless, these antigens successfully presented the mAb806-epitope in its antibody-binding conformation, and may thus offer a useful starting point for further development. Modifications of both the immunogen composition^33–35^ and immunization protocols,^36,37^ in order to enhance immunogenicity and focus the antibody response against the epitope,^11^ could potentially lead to immunogens that can overcome the obstacles outlined above and robustly elicit mAb806-like, cancer-selective antibodies.

## Methods

### Molecular dynamics simulations

For the starting structures of our EGFR simulations (simulations 1, 2, 5, and 6), four frames were selected randomly from an equilibration simulation of an untethered EGFR ectodomain crystal structure (PDB ID: 3NJP); the double mutation C271A/C283A was introduced for simulations 5 and 6. The CR2 domain (also known as domain IV; residues Val481 through Thr614) was omitted from the ectodomain in these simulations to reduce the size of the molecular system and accelerate conformational dynamics. mAb806 was docked onto EGFR in frames selected at a regular time interval from simulations 1, 2, 5, and 6, by aligning the Cα atoms of the epitope in the crystal structure of the mAb806-epitope complex (PDB ID: 3G5V) onto the epitope in the simulated EGFR conformation. We then used selected frames from simulation 2 to generate a structural model of the mAb806-EGFR complex, from which we initiated two additional simulations (simulations 3 and 4). In all six of these simulations, the DES-Amber SF1.0 force field^38^ was used for the protein, together with the TIP4P-D water model.^39^ Simulations 1, 2, 3, 5, and 6 used Times Square sampling^40^ (an adaptive variant of simulated tempering^41,42^) so that the systems traversed a temperature range between 310 K and 350 K, thereby accelerating sampling of the protein conformations; simulation 4 was run at a constant temperature of 310 K. All systems were maintained at a pressure of 1 atm.

Isobaric-isothermal simulations for the free epitope peptide and each of the designed antigens were performed at a temperature of 310 K and a pressure of 1 atm, using three different force fields: 1) amber99SB*-ILDN^43^ (which builds on other modifications^44,45^ to Amber99^46^) with the TIP3P water model,^47^ 2) DES-Amber SF1.0 with the TIP4P-D water model, and 3) Charmm22*^48^ with the TIP3P-CHARMM water model.^49^

In all simulations, bond lengths to hydrogen atoms were constrained using an implementation^50^ of M-SHAKE,^51^ electrostatic interactions were computed using the *u*-series method,^52^ and Lennard-Jones interactions were calculated using a cutoff of 12 Å. The pressure and temperature were controlled using the Martyna-Tobias-Klein (MTK) algorithm^53^ and the Nosé-Hoover thermostat,^54^ respectively, applied using the multigrator framework.^55^ The RESPA multiple time scale integrator^56^ was used to propagate the dynamics, updating the bonded, Lennard-Jones, and short-range electrostatic forces every 2.5 fs, and the long range electrostatic forces every 5 fs. All simulations were performed on the Anton 2 specialized supercomputer.^57^

### Expression and purification of DKK1-6 and DKK1-16

DKK1-6 and DKK1-16 constructs, including an N-terminal His6-SUMO tag, were expressed in Sf21 insect cells grown in Insect-XPRESS medium (Lonza) at 27 °C for 66 hours. To facilitate disulfide bond formation, the proteins were secreted in the medium. After clarification of the medium by centrifugation, the proteins were isolated using HisPur Ni-NTA resin (ThermoFisher) equilibrated in buffer A (20 mM Tris/HCl, 500 mM NaCl, 20 mM imidazole, pH 7.1). The resin was washed extensively with buffer A, followed by elution with buffer B (20 mM Tris/HCl, 500 mM NaCl, 500 mM imidazole, pH 7.1). Subsequently, the proteins were dialyzed into buffer C (20 mM Tris/HCl, 50 mM NaCl, pH 7.1) in the presence of Ulp1 protease, in order to cleave the His6-SUMO tag. The cleaved proteins were further purified by cation exchange chromatography using HiTrap SP HP column (GE Healthcare) equilibrated in buffer C. After washing with buffer C, the proteins were eluted by applying a linear gradient from 0 to 100% of buffer D (20 mM Tris/HCl, 1 M NaCl, pH 7.1) in 30 column volumes. Elution fractions containing pure DKK1-6 and DKK1-16, as determined by SDS-PAGE, were deglycosylated by incubation with PNGase F (New England BioLabs) at 37 °C for one hour. Deglycosylated proteins were purified to homogeneity by size exclusion chromatography using a HiLoad Superdex 75 PG column (GE Healthcare) equilibrated in Dulbecco’s Phosphate buffered saline (DPBS; Gibco). Subsequently, endotoxins were removed using Pierce High-Capacity Endotoxin Removal Resin (Thermo Scientific). The proteins were flash frozen in liquid nitrogen and stored at a concentration of approximately 10 mg/mL at −80 °C.

### Stable cell lines

Stable cell lines of U87MG^wt^ ^EGFR^ and U87MG^EGFRvIII^ were produced by Genscript under a research contract. Briefly, the full-length cDNA of human EGFR and mutant EGFRvIII were synthesized and subcloned into the pLenti-puro vector, and the recombinant lentivirus was packaged in 293T cells. U87MG cells were transduced with the recombinant lentivirus, centrifuged at 800 *g* for 120 minutes at 20 °C, incubated for 24 hours, and then cultured in fresh complete growth medium for an additional 24 hours. The transduced cells were selected for four days in medium containing 0.75 μg/mL (for U87MG^wt^ ^EGFR^) or 1 μg/mL (for U87MG^EGFRvIII^) puromycin. Elevated expression levels of EGFR and EGFRvIII were confirmed by quantitative polymerase chain reaction (qPCR) and by western blot analysis (Figure S6). U87MG, A431, and A549 cell lines were obtained from ATCC.

### Surface plasmon resonance

Surface plasmon resonance (SPR) binding data were collected on a ProteOn XPR36 biosensor using GLC sensor chips. Antibody ch806 was captured onto a Protein A–coated sensor at five different surface densities. All studies were run in 10 mM HEPES, 150 mM NaCl, 0.005% tween-20, 3.4 mM EDTA, and 0.2 mg/mL BSA at pH 7.4 and 25 °C. The epitope peptide, DKK1-6, and DKK1-16 were tested in a three-fold concentration series up to the highest concentrations of 4 μM, 1 μM, and 333 nM, respectively. Interspot referenced response data from five different surface densities of antibodies were globally fit to a 1:1 binding stoichiometry to extract estimates of the binding rate constants.

### Microscale thermophoresis

The fluorescent label AlexFluor-647 (ThermoFisher) was covalently attached by N-hydroxysuccinimide (NHS) coupling to the N-terminal primary amine of the ch806 antibody. The experiments were performed on the Monolith NT.115 instrument using the RED detector, with the ch806 concentration at 70 nM. The epitope peptide was tested in a 12-point, three-fold serial dilution beginning at a 1000 μM peptide concentration, whereas DKK1-6 and DKK1-16 were tested in a 12-point, two-fold serial dilution beginning at 72.5 μM and 54.5 μM, respectively. The buffer used in the assays contained 20 mM HEPES and 150 mM NaCl, with pH 7.5. After a 30-minute incubation, the samples were loaded into MST NT.115 standard glass capillaries and a microscale thermophoresis (MST) analysis was performed using 60% LED power and 40% MST power. At high concentrations, analytes can stick to the capillary walls, leading to artificial changes in the thermophoresis. We thus omitted the data points at high concentrations (>111 μM for the epitope peptide, >9.4 μM for DKK1-6, and >6.7 μM for DKK1-16) from our estimations of the binding affinities.

### One-dimensional NMR spectroscopy

Proteins were measured in DPBS buffer with the addition of 10% D_2_O. Proton 1D spectra were acquired at 25 °C. Water suppression sequence (Watergate) was used to eliminate water resonance at 4.7 ppm. 128 scans were recorded for each spectrum. For TCEP-reduced samples, the proteins were incubated with 2 mM TCEP at room temperature for 20 minutes prior to measurements.

### Immunization of rabbits

Two New Zealand rabbits were immunized with each immunogen. 1 mL of serum from each rabbit was collected via ear vein three days before the first immunization (prebleed). In the first immunization, 200 μg of immunogen, mixed with Complete Freund’s Adjuvant (CFA), was injected subcutaneously at four spots on the rabbit’s back. In subsequent immunizations, 200 μg of immunogen, mixed with Incomplete Freund’s Adjuvant (IFA), was similarly injected. 100 μL of anti-sera were drawn from each rabbit via ear vein one week after the second and third immunizations for tests of antigen binding by ELISA. Polyclonal antibodies were affinity-purified from the 10–15 mL of anti-sera drawn one week after the fourth immunization; the anti-sera from rabbits immunized by the designed antigens were purified using the corresponding protein antigens coupled to a Sepharose 4B column, whereas the anti-sera from rabbits immunized by epitope-peptide KLH-conjugate were purified using the epitope peptide [Cyclo(5,20)] H_2_N-SGGGCGADSYEMEEDGVRKC-OH coupled to a Sepharose 4B column. The antibody concentrations were determined by absorbance at 280 nm with a NanoDrop 2000 spectrophotometer.

### Enzyme-linked immunosorbent assay (ELISA)

ELISA plates were coated by incubating with antigen solutions (pH 7.4) at a concentration range of 1–10 μg/mL for two hours at 37 °C or overnight at 4 °C. The plates were then washed three times with 200 μL of washing buffer (PBS buffer containing 0.05% Tween 20), and incubated with 200 μL blocking buffer (PBS buffer containing 0.05% Tween 20 and 0.01 g/mL bovine serum albumin) for two hours at 37 °C to block non-specific binding sites in the coated wells.

The anti-sera or purified polyclonal antibodies were diluted with the above blocking buffer, 100 μL of the solution was added to each well, and the plate was incubated for one hour at 37 °C or overnight at 4 °C. The plate was washed three times with 200 μL of washing buffer.

Horseradish peroxidase-conjugated anti-rabbit IgG antibody was diluted in the blocking buffer, 100 μL of the solution was added to each well, and the plate was incubated for 30 minutes at 37 °C. The plate was washed five times with 200 μL of washing buffer. 100 μL of TMB reagent was added to each well, and after sufficient color development (15–30 minutes), 100 μL of stopping buffer (1 M HCl) was added to each well. The absorbance was measured at 450 nm with a plate reader.

## Acknowledgments

We thank David Borhani and Michael Eastwood for helpful discussions and critical readings of the manuscript, Jordan Caleb and Kevin Yuh for preparing the molecular graphics, Jordan Caleb for preparing the supplementary movie, and Jessica McGillen and Eric Martens for editorial assistance.

## Notes

### Competing Interest Statement

The authors have declared no competing interest.

