## Supplementary Information for "Design of antigens to present a tumor-specific cryptic epitope"

### Supplementary Information:
### Design of antigens to present a tumor-specific cryptic epitope

#### Supplementary Methods

##### ***Computer-aided design of antigens***

###### *Generation of conformational ensemble of the Fab-epitope complex*

A molecular dynamics (MD) simulation of the complex between mAb806 and EGFRΔCR2 (the ectodomain of EGFR with domain IV deleted) was carried out for 10 μs at 310 K and 1 atm, starting from a structural model that was generated by superimposing the mAb806-epitope crystal structure (PDB ID: 3G5V) onto a simulated conformation of the apo EGFRΔCR2, aligning the Cα atoms of the epitope. 100 snapshots were taken at 100-ns intervals, and the Fab and epitope (residues C287–C302 in EGFR) conformations were extracted from these snapshots. These conformations were used in the next step of searching for compatible scaffolds onto which the epitope could be grafted.

###### *Computational search for scaffolds suitable for grafting the epitope*

The motif graft protocol in the Rosetta Protein Modeling software was used to search a set of 21,829 PDB structures for scaffolds onto which the epitope could be grafted. A scaffold was considered to be a candidate if the following criteria were met: 1) the epitope could be superimposed onto a replacement segment of the same length in the scaffold with a Cα root-mean-square deviation (RMSD) no greater than 1.5 Å; 2) the N, Cα, C RMSD between the N- and C-terminal residues in the epitope and those in the replacement segment was no greater than 1.5 Å; and 3) when the scaffold's residues were mutated to Gly and the replacement segment was excluded, at most 20 atoms in the scaffold sterically clashed with the superimposed epitope-antibody complex. The replacement segment in the candidate scaffold was then replaced by the epitope sequence, and these grafts were used in the next step of computational and manual redesign.

*Computational redesign of grafts*

Each scaffold protein with the grafted epitope underwent an additional round of computational design: We used the RosettaRemodel program[1] in the Rosetta software package to select stabilizing mutations in residues that were in the vicinity of the epitope (including in the epitope itself), while maintaining the key interactions between the epitope and the antibody (by preserving the amino acids C287, E293, and E295–C302). The designed proteins were subjected to the next step of filtering according to the results of MD simulations.

*Filtering of the designs informed by MD simulations*

A molecular dynamics simulation of each designed protein was carried out at 310 K and 1 atm for a duration between 10 and 100 μs (see Methods in main text for additional information). A protein was eliminated from further consideration if its epitope substantially deviated from the antibody binding conformation during its simulation.

***On-Cell Western assay***

The binding of antibodies to cells was measured by an on-cell Western assay. The cells from T75 culture flasks were trypsinized using 0.025% Trypsin-EDTA and washed twice with fresh media. The cells were then distributed into 96-well, poly-D-lysine–treated culture plates (Corning), which were opaque black with clear bottoms, at 50,000 cells per well using appropriate media for each cell line. The cells were incubated at 37 °C overnight to allow for the recovery of receptors and adherence, then washed three times with cold PBS buffer and blocked with 50 μL blocking buffer (Becton Dickinson human Fc block) for 0.5 hours on ice. 50 μL of an antibody of interest or the human IgG1 isotype control (Biolegend) was added to the cells in a series of three-fold dilutions of the blocking buffer, and the cells were incubated for two hours at 4 °C. The cells were then washed three times with cold PBS. Anti-human IgG secondary antibody conjugated with IRDye 800CW at a 1:400 ratio to the primary antibodies, and DRAQ5 diluted in the blocking buffer, were added to the cells, and the cells were incubated for one hour at 4 °C. The cells were then washed three times with cold PBS, and air dried. The fluorescence (F) at 700 nm (DRAQ5) and 800 nm (secondary antibody) was read on Li-Cor Odyssey. The specific normalized signal was calculated according to the equation:

$$\text{relative binding} = \frac{F_{800nm,\text{with antibody}} - F_{800nm,\text{without antibody}}}{F_{700nm}}$$

The binding of IgGs to cells was analyzed as follows. The binding is bivalent:

$$\mathrm{Ab} + \mathrm{Ag} \overset{K_D}{\rightleftharpoons} \mathrm{Ab \cdot Ag}$$

$$\mathrm{Ab \cdot Ag} + \mathrm{Ag} \overset{\gamma\, K_D}{\rightleftharpoons} \mathrm{Ab \cdot Ag_2}$$

Here, Ab denotes the antibody (IgG), and Ag the antigen. $K_D$ is the equilibrium dissociation constant for monovalent binding between one Fab of the IgG and the antigen. The factor $\gamma$ quantifies the local concentration enhancement for binding to the second surface antigen due to anchoring by binding to the first antigen. It is in units of length and is determined by the relative positions of the two Fabs in the same IgG molecule.[2]

At equilibrium, the molecular concentrations satisfy the following:

$$\frac{[\mathrm{Ab}][\mathrm{Ag}]}{[\mathrm{Ab}\cdot\mathrm{Ag}]} = K_D$$

$$\frac{[\mathrm{Ab}\cdot\mathrm{Ag}][\mathrm{Ag}]}{[\mathrm{Ab}\cdot\mathrm{Ag}_2]} = \gamma K_D$$

$$[\mathrm{Ab}] + \frac{n\cdot S}{V}[\mathrm{Ab}\cdot\mathrm{Ag}] + \frac{n\cdot S}{V}[\mathrm{Ab}\cdot\mathrm{Ag}_2] = C_{\mathrm{Ab}}$$

$$[\mathrm{Ag}] + [\mathrm{Ab}\cdot\mathrm{Ag}] + 2[\mathrm{Ab}\cdot\mathrm{Ag}_2] = C_{\mathrm{Ag}}$$

Note that [Ab] is the bulk density of the antibody, but [Ag], [Ab · Ag], and $[\mathrm{Ab}\cdot\mathrm{Ag}_2]$ are surface densities on the cell. Here $S$ is the average cell surface area, $n$ is the number of cells in the solution, and $V$ is the total volume of the solution. (In our experiments, $n$ = 50,000 cells are suspended in a solution of $V$ = 100 μL; assuming an average cell diameter of 25 μm, we have $\phi \equiv nS/V = 9.8 \times 10^{-4}\ \mu\mathrm{m}^{-1}$.) $C_{\mathrm{Ab}}$ is the total bulk density of the antibody, and $C_{\mathrm{Ag}}$ is the average total surface density of the antigen.

The relative binding measured by the on-cell Western assay is given by:

$$\text{relative binding} = \chi \cdot ([\mathrm{Ab}\cdot\mathrm{Ag}] + [\mathrm{Ab}\cdot\mathrm{Ag}_2])$$

where $\chi$ is a constant across all binding experiments. The preceding equations allowed us to determine the unknown variables, $K_D$, $\gamma$, and $C_{\mathrm{Ag}}$, by fitting to the measured relative binding at different antibody concentrations (i.e., different $C_{\mathrm{Ab}}$) and against different cell lines.

This fitting enabled us to determine the local concentration enhancement, $\gamma$, and three equilibrium dissociation constants: $K_{D,\mathrm{cetuximab}}$ for binding of cetuximab to EGFR, $K_{D,\mathrm{ch806}}$ for binding of ch806 to (misfolded) EGFR, and $K_{D,\mathrm{vIII}}$ for binding of ch806 to EGFRvIII. We also determined the surface density of $C_{\mathrm{Ag}}$ binding-competent EGFR on different cells. The surface densities of binding-competent EGFR on the same cell were different for cetuximab and ch806, since the latter presumably bound only to the misfolded subpopulation.

#### Supplementary Figures

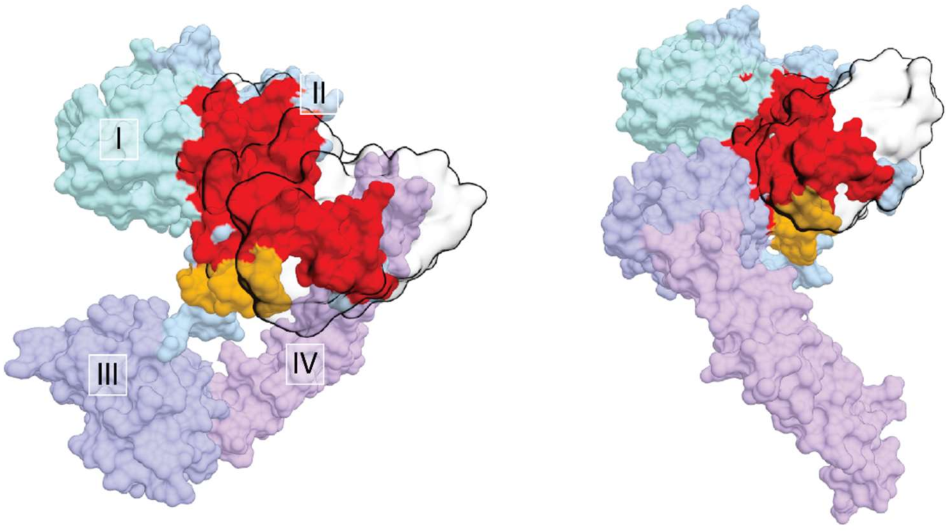


**Figure S1**. The epitope is inaccessible to mAb806 in known crystal structures of EGFR. Substantial steric clashes would occur if mAb806 were docked onto the epitope in either the tethered (auto-inhibited) or untethered (active) conformation of EGFR. Here, mAb806 was docked onto the two crystal structures of EGFR by aligning the C$\alpha$ atoms of the epitope in the crystal structure of the mAb806-epitope complex (PDB ID: 3G5V) onto the epitope in the crystal structures of the tethered (PDB ID: 1NQL) and untethered (PDB ID: 3NJP) conformations. The antibody is depicted as hollow contours. Domains I (turquoise), II (blue), III (purple), and IV (lavender) of EGFR are shown in molecular surface representation, as is the epitope (orange), which is part of domain II. The EGFR atoms in steric collision with the docked mAb806 (inter-atomic distance ≤ 1.5 Å) are highlighted in red.

A

B

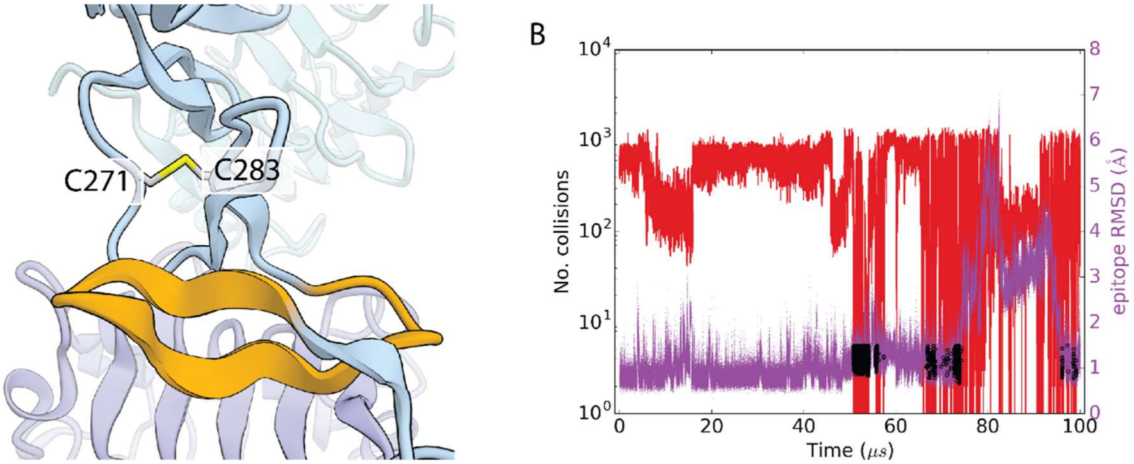


**Figure S2**. Breaking a disulfide bond between residues C271 and C283 near the epitope favors local unfolding and increases the occurrence of mAb806-binding conformations. **A**) The disulfide bond between residues C271 and C283 (yellow) helps to maintain the native conformation of domain II, excluding the epitope (orange) from mAb806 binding. The double mutant C271A, C283A eliminates this disulfide bond. **B**) In our MD simulations with simulated tempering (simulations 5 and 6; only simulation 5 is shown here), the double mutant presented the epitope in the binding-competent conformation—highlighted by black circles—much more frequently than the wild-type EGFR (compare Fig 2C), in agreement with the experimental observation[5] that mAb806 binds to this double mutant with much higher affinity than to the wild type.

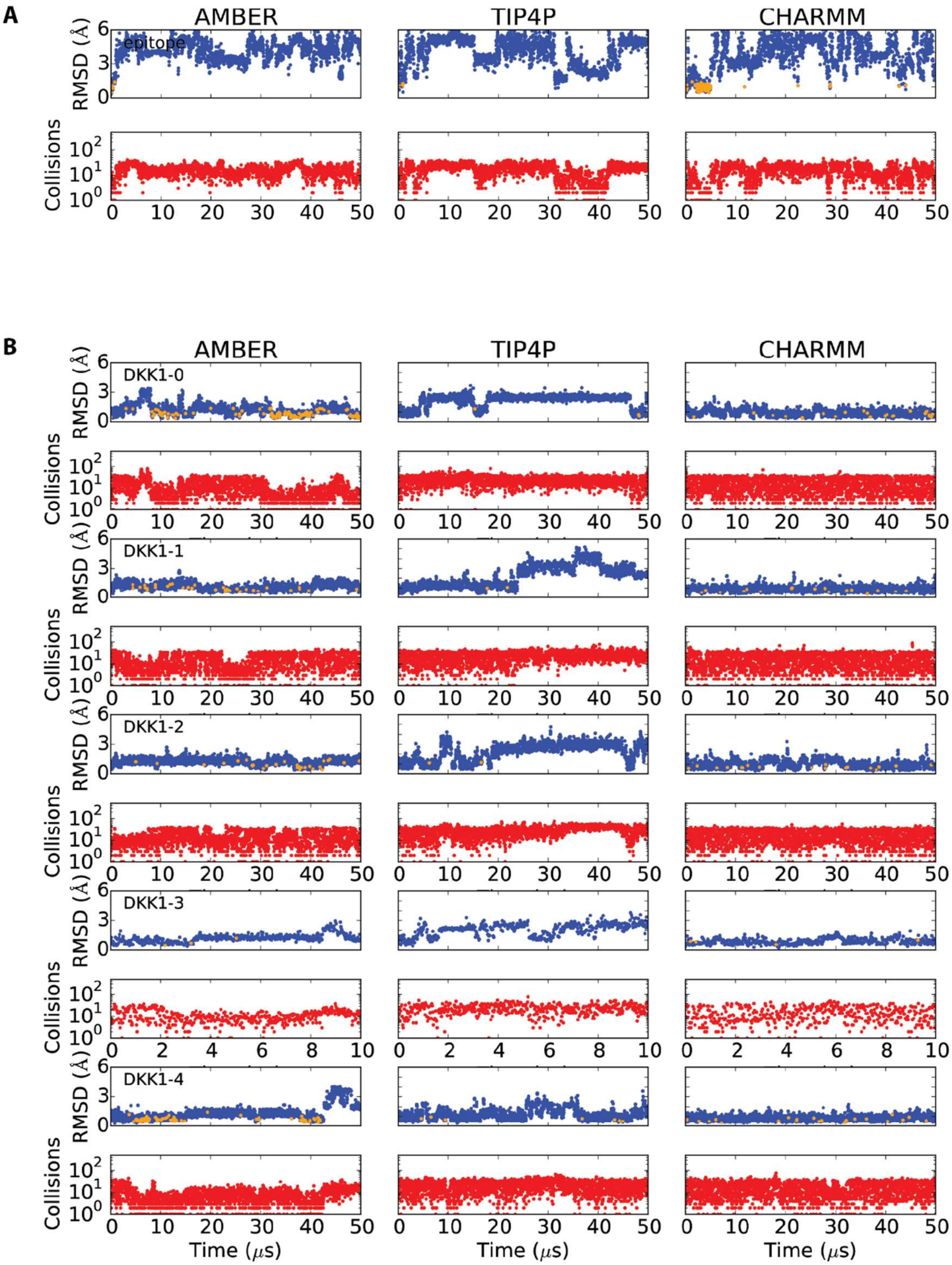


(figure continued on next page)

**B** (page 2)

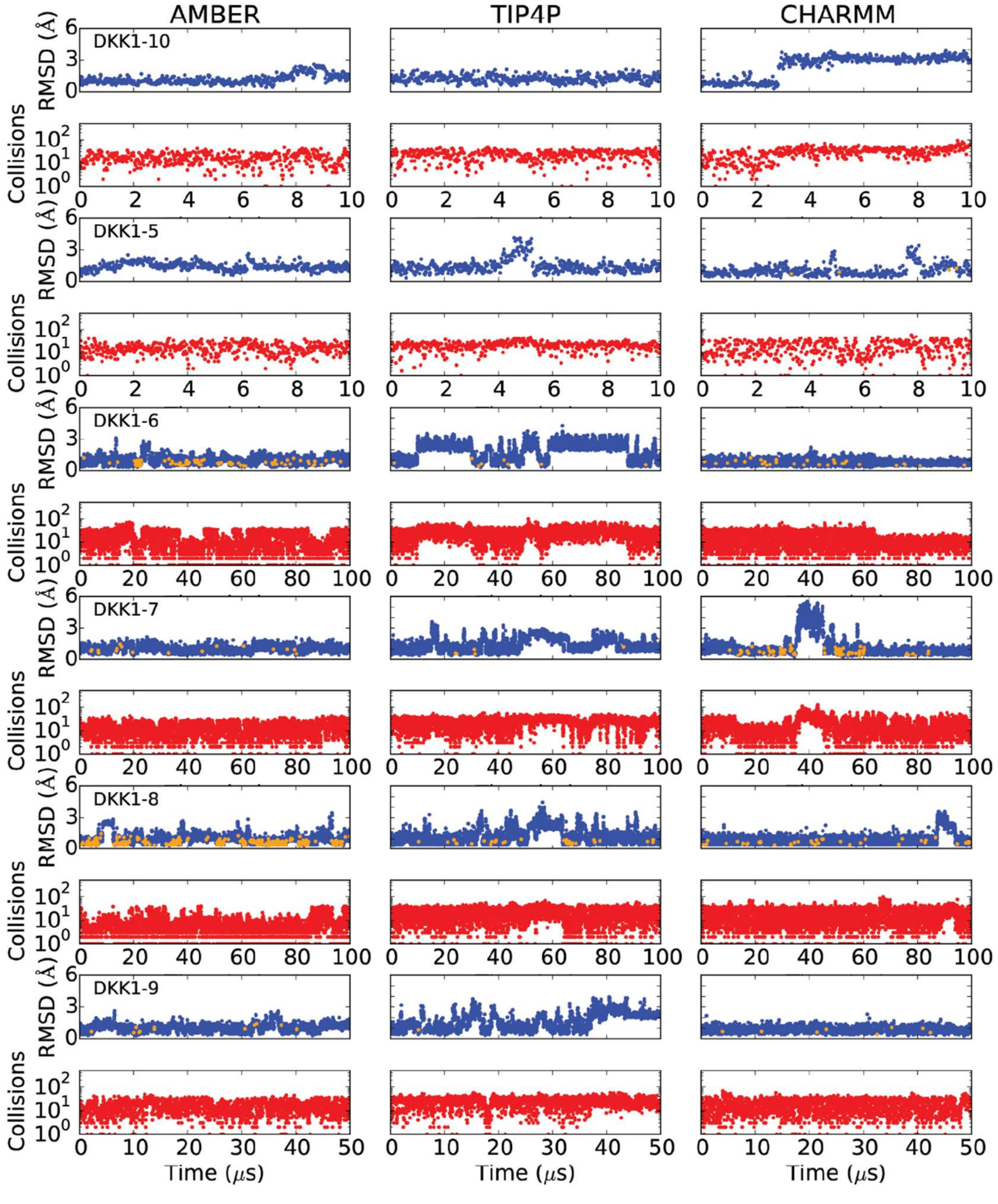


(figure continued on next page)

**B** (page 3)

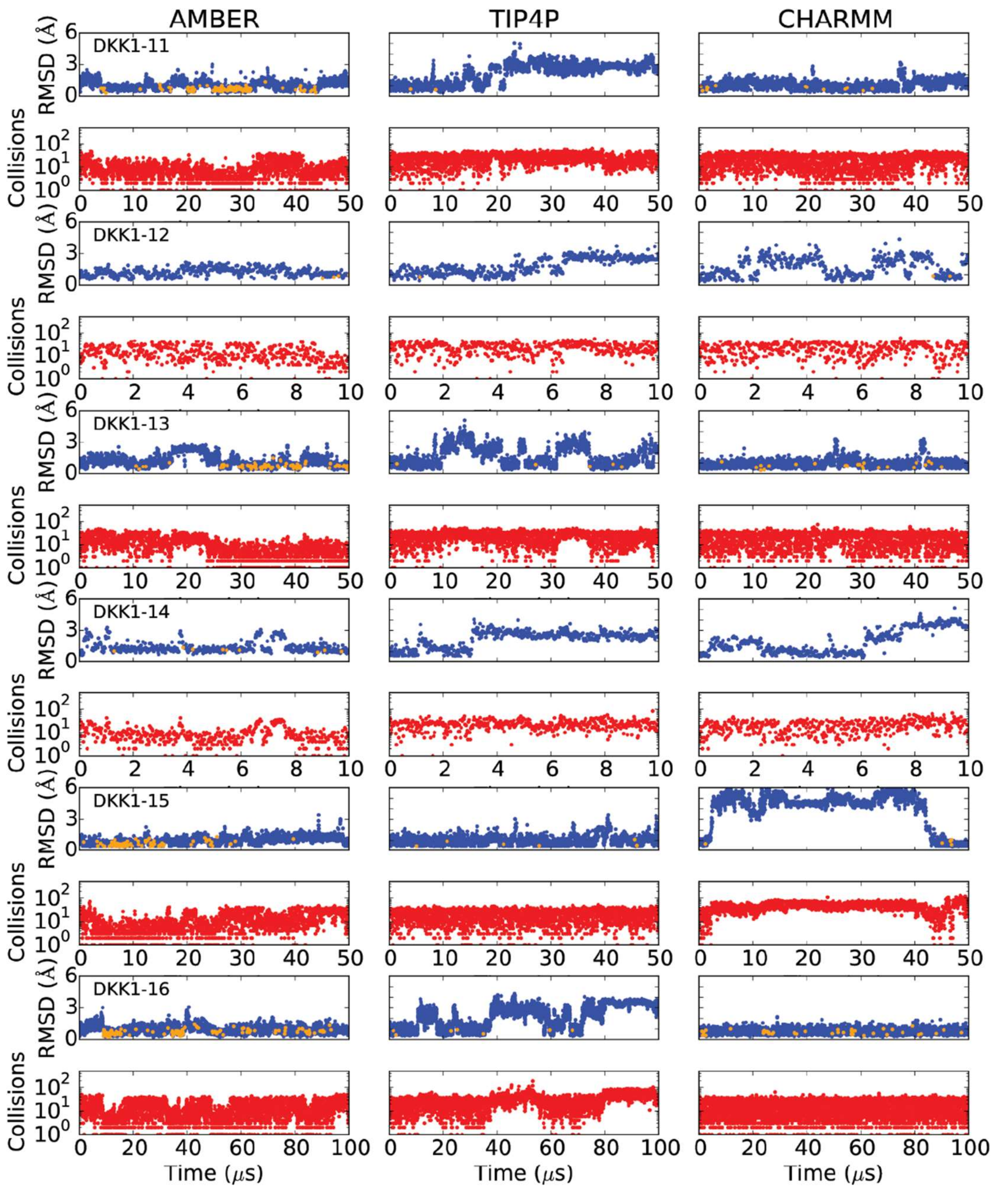


**Figure S3**. Sampling of the mAb806-binding conformation by the epitope in **A**) the stand-alone

16-mer peptide and **B**) the designed antigens, in our MD simulations using three force fields (AMBER: Amber99SB*-ILDN + TIP3P water model; TIP4P: DES-Amber SF1.0 + TIP4P-D water model; CHARMM: CHARM22* + TIP3P-CHARMM water model). The Cα RMSD from the mAb806-binding conformation in the crystal structure of the mAb806-epitope peptide complex (PDB ID: 3G5V) is plotted in blue in the odd-numbered rows, and the number of atoms in potential steric collision with the antibody when the peptide or designed antigen is docked onto mAb806 is plotted in red in the even-numbered rows. The orange points in the RMSD plots indicate frames in which the RMSD is no greater than 1.5 Å and there are no colliding atoms. In contrast to the stand-alone peptide, which was unstructured and rarely adopted the mAb806-binding conformation in our simulations, the epitope in most of the designed antigens retained or predominantly occupied the mAb806-binding conformation.

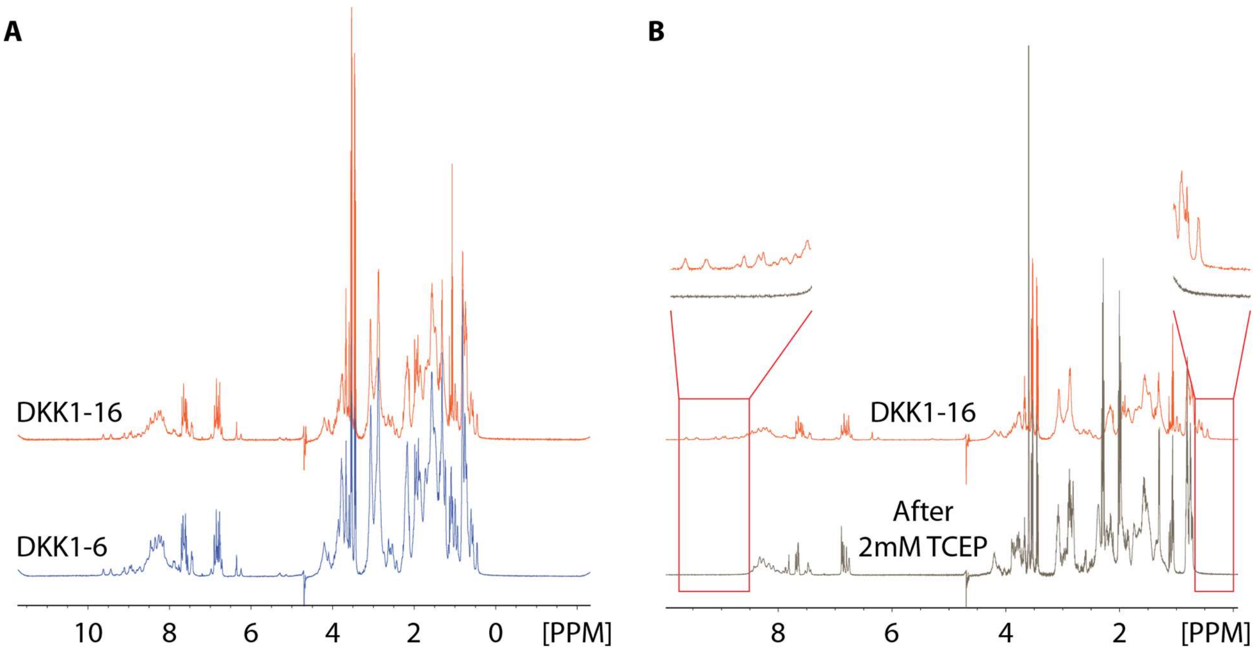


**Figure S4**. Structural characterization of the designed antigens. **A**) The designed antigens DKK1-6 (blue trace) and DKK1-16 (red trace) adopted similar folded conformations, as shown by 1D NMR spectroscopy. Both designs exhibit the signatures of folded proteins, showing dispersion of resonances in 0–0.8 ppm (aliphatic) and 8.4–9.6 ppm (amide/aromatic) intervals. There are also unstructured regions in the two proteins, probably in the long loop near the C-terminus. **B**) After incubation with 2 mM TCEP for 20 minutes at room temperature (black trace), which reduced the disulfide bonds, the dispersion peaks disappeared (highlighted by the magnified regions, in pink boxes), indicating that the protein unfolded. This suggests that the disulfide bonds are indispensable to the structural integrity of the designed proteins.

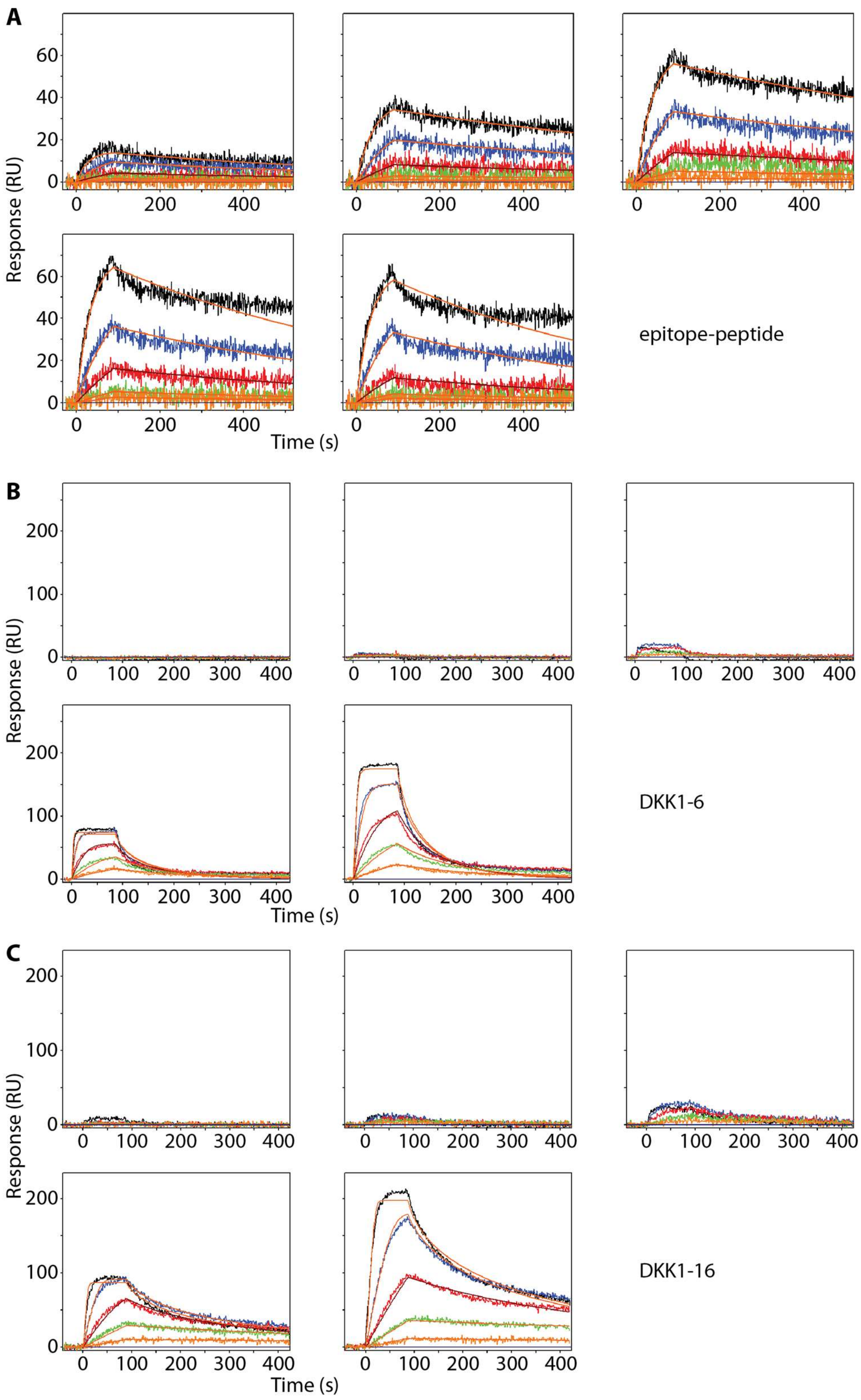


**Figure S5**. Surface plasmon resonance data for the binding of the ch806 antibody to **A**) the epitope peptide, **B**) DKK1-6, and **C**) DKK1-16. In each panel, the series of five plots corresponds to increasing relative surface densities of the ch806 antibody, and the time traces in each plot show antigen concentrations in three-fold dilution series from a maximum concentration of 4 μM (for A, the epitope peptide), 1 μM (for B, DKK1-6), and 333 nM (for C, DKK1-16), with black denoting the maximum concentration and blue, red, green, and orange denoting the successive dilutions. The overlaid curves represent best global fits used to derive binding rate constants (Methods). In each time trace, the initial peak indicates binding of the antibody to the antigen, and the subsequent decline indicates dissociation of the antibody from the antigen. RU, resonance units.

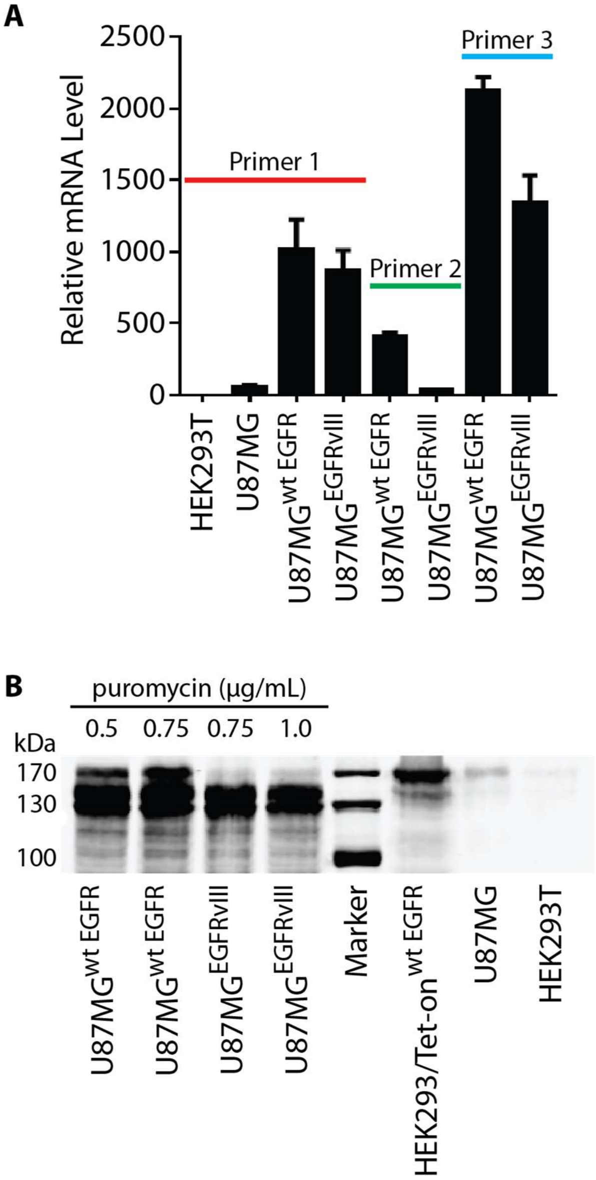


**Figure S6**. Expression of EGFR and EGFRvIII in the engineered stable U87MG$^{wt\ EGFR}$ and U87MG$^{EGFRvIII}$ cells. **A**) Quantitative polymerase chain reaction (qPCR) measurements of relative mRNA levels in the HEK293T cells, parental U87MG cells, and engineered stable cell lines. Primer 1 (forward: CCTGGTCTGGAAGTACGCAG, reverse: GCGATGGACGGGATCTTAGG) and primer 3 (forward: GCTATGAGATGGAGGAAGACG, reverse: TCACCAATACCTATTCCGTTACAC) amplify both wild-type EGFR and EGFRvIII, whereas primer 2 (forward: GCCTCCAGAGGATGTTCAAT, reverse: GACATAACCAGCCACCTCCT) amplifies only wild-type EGFR but not EGFRvIII. The mRNA levels of EGFR and EGFRvIII in the engineered cell lines are ~15 and ~13 times, respectively, that of the EGFR mRNA in the parental U87MG cells. RNA was isolated from the cells using the Trizol extraction kit. qPCR was performed with one minute pre-denaturation at 95 °C, followed by 40 cycles of denaturation at 95 °C for 15 seconds, annealing at 60 °C for 15 seconds, and extension at 72 °C for 45 seconds. The fluorescence readout was collected in the extension step. Each sample was measured in quadruplicate. **B**) Western blot of cell lysates. The engineered cells were selected by resistance to the antibiotic puromycin at the indicated concentrations. The EGFR and EGFRvIII were detected using the rabbit monoclonal antibody against EGFR (Cell Signaling Technology, Catalog # 4267) with a goat anti-rabbit IgG secondary antibody (Li-COR, Catalog # 926-32211). Different bands may correspond to different glycosylation states (EGFR has 16 potential N-glycosylation sites and EGFRvIII has 12).

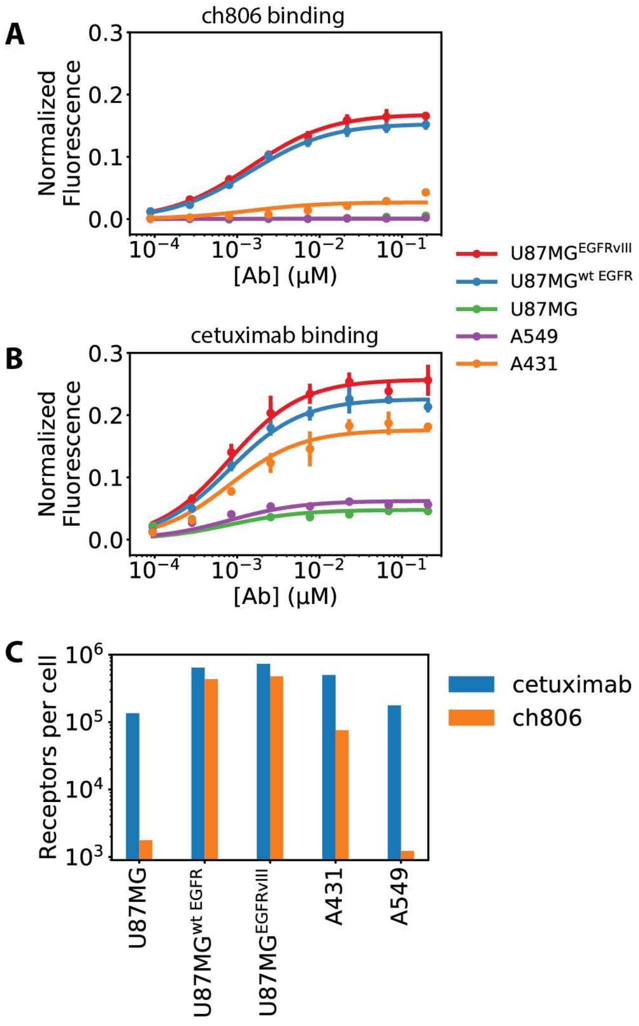


**Figure S7**. Binding of monoclonal antibodies ch806 and cetuximab to various tumor cell lines, as measured by an On-cell Western assay. **A**) ch806 binds only to tumor cell lines that either overexpress EGFR (A431 and U87MG$^{wt\ EGFR}$) or express the EGFRvIII mutant (U87MG$^{EGFRvIII}$). Shown are the experimental results (filled circles) and theoretical binding curves (solid curves) fitted to the data. The normalized fluorescence is the ratio of fluorescence from the surface-bound secondary antibodies to the fluorescence from DNA-bound fluorophores, and quantifies the EGFR-bound antibodies per cell (see Supplementary Methods). **B**) Cetuximab binds to all five cell lines tested. **C**) Number of surface receptors on each cell line for each antibody.

Compared to cetuximab, ch806 binds to a subpopulation—presumably the misfolded portion—of surface EGFR. The number of surface receptors and equilibrium binding constants are determined by simultaneously fitting the binding equations to all of the binding data, assuming that there are $5 \times 10^5$ EGFR molecules on each A431 cell.[3] This yields $K_{D,\mathrm{ch806}}$ = 1.4 nM and $K_{D,\mathrm{cetuximab}}$ = 0.8 nM. The $K_D$ for cetuximab is in good agreement with previous measurements of cetuximab binding to EGFR (2.3 nM[4] and 0.53 nM[3]), validating our On-cell Western assay for quantitative characterization of antibody binding to cell surface receptors.

#### Supplementary Movies

**Movie S1**. Conformational dynamics of the EGFR ectodomain in simulation 2 (an analysis of which is shown in Figure 2). The simulation was initiated from the untethered conformation. **0–9 seconds of the movie**: Docking MAb806 Fab (purple) onto its epitope results in substantial steric collisions (the part of the Fab in collision is highlighted in red). The proteins are shown in the cartoon representation, with the epitope highlighted in orange and the rest of EGFR in gray. **10–25 seconds**: The untethered (light blue) and tethered (green) conformations are superimposed onto the initial conformation of the MD simulation by aligning their L1 domains. Drastic differences in the positions of their L2 and CR2 domains between the untethered and tethered conformations are emphasized in red. **26–27 seconds**: We omitted domain IV (the CR2 domain) of the EGFR ectodomain from our simulation, as shown by the comparison between the simulated construct and the full EGFR ectodomain in untethered conformation. **28–48 seconds**: The EGFR ectodomain can transition from the untethered to tethered conformation while hiding the epitope from mAb806-binding. The color of the epitope indicates the number of atoms in steric collision with a docked mAb806 Fab: the redder the color, the more number of atoms in collision; green indicates zero collisions. The good alignment to the crystal structure of the tethered conformation (green) at the end of this simulation segment shows that the EGFR has transitioned from the untethered to tethered conformation. **49–67 seconds**: The EGFR ectodomain, after sampling a wide range of conformations, transiently exposes the epitope, to which the mAb806 Fab (purple) can be docked without steric collision. **68–86 seconds**: The EGFR ectodomain samples additional conformations that expose the epitope to mAb806 binding. Each such conformation is retained in the background in faded colors. **87–89 seconds**: All the mAb806-binding conformations of EGFR sampled in the simulation are shown, together with the crystal structure (purple) of mAb806 Fab in complex with the epitope peptide.  All structures are aligned by the Cα atoms in the epitope.
